# Dose-dependent myelin g-ratio alterations in the white matter of 16p11.2 copy number variations

**DOI:** 10.64898/2026.10.19.760158

**Authors:** Wen Da Lu, Charles-Olivier Martin, Khadije Jizi, Mark C. Nelson, Bogdan Draganski, Sebastien Jacquemont, Christine L. Tardif

## Abstract

Copy number variations (CNVs) at the 16p11.2 locus are major genetic risk factors for neurodevelopmental conditions and drive reciprocal alterations in cerebral white matter structure and connectivity. However, whether these alterations reflect differences in the axonal or myelin compartments remains unresolved with conventional diffusion MRI. We combined diffusion MRI with magnetization transfer saturation mapping across two ComBat-harmonized cohorts comprising 80 controls, 26 16p11.2 deletion carriers, and 13 duplication carriers. We quantified axonal volume, myelin volume, and aggregate g-ratio of white matter tracts forming the structural connectome. Metrics were evaluated using normative modeling across tract length, resting-state functional networks, and cortical nodes. We additionally examined 29 individuals carrying other neurodevelopmental CNVs. 16p11.2 gene dosage was associated with regional reciprocal differences in tract axonal volume, with greater volume in deletion carriers and lower volume in duplication carriers. In contrast, myelin volume did not differ significantly from controls. This mismatch resulted in pervasive aggregate g-ratio alterations, with higher g-ratios (relatively thinner myelin sheaths) in deletion carriers and lower g-ratios (relatively thicker sheaths) in duplication carriers. Regional g-ratio alterations also extended to other neurodevelopmental CNVs. These findings resolve some of the biological ambiguity underlying white matter alterations in 16p11.2 CNV carriers, implicating the axonal compartment while revealing comparatively stable myelin volume. The disrupted axon-myelin scaling results in connectome-wide g-ratio alterations and may represent a shared microstructural hallmark of genetic neurodevelopmental risk.

## Introduction

Copy number variations (CNVs) at the 16p11.2 locus represent one of the most prevalent and highly penetrant genetic risk factors for neurodevelopmental and psychiatric conditions.^1^ Deletion and duplication carriers exhibit contrasting physical profiles, notably in head circumference (macrocephaly vs. microcephaly), platelet count, and adiposity,^2–4^ alongside shared vulnerabilities to intellectual disability, attention-deficit/hyperactivity disorder, and autism spectrum disorder (present in 16-34% of carriers).^5^ However, other psychiatric and cognitive trajectories diverge significantly: duplication carriers face a 10-fold higher risk of developing schizophrenia,^4^ whereas deletion carriers are uniquely characterized by severe impairments in phonology.^3^

These opposing clinical phenotypes are mirrored by reciprocal alterations in macroscopic brain structure and function. Structural MRI studies demonstrate reciprocal differences in brain volume and cortical surface area, characterized by an increase in deletion carries and widespread reduction in duplication carriers,^6^ as well as bidirectional thickness changes in the corpus callosum.^7^ Beyond structural anatomy, resting-state functional MRI study revealed a connectome-wide gene dosage mirror effect in 16p11.2 carriers at both the global connectivity and network levels.^8^

A reciprocal gene-dosage effect has also been reported in the microstructure of white matter tracts of 16p11.2 CNV carriers using diffusion-weighted MRI (dMRI). These findings are primarily based on conventional diffusion tensor metrics, including fractional anisotropy (FA) and axial diffusivity (AD), which provide sensitive but biologically nonspecific markers of white matter microstructure. FA and AD were elevated in deletion carriers and reduced in duplication carriers across specific regions.^9–11^ Although highly sensitive, these measures are biologically nonspecific, conflating differences in axonal density, myelination, and fiber geometry. Resolving this biological ambiguity is critical, as axonal versus myelin alterations carry vastly different developmental implications and would require distinct therapeutic interventions.

Recent work utilizing advanced dMRI acquisitions and multi-compartment microstructural modeling has begun to bridge this gap, identifying significant increases in the total intra-axonal volume of white matter tracts in children (2–8 years old) with 16p11.2 deletions.^12^ However, this technique is insensitive to the myelin compartment because the short T2 relaxation time of water within the myelin sheath precludes its detection with conventional dMRI,^13^ leaving white matter myelination uncharacterized in 16p11.2. The thickness of the myelin sheath relative to the axon caliber can be quantified using the g-ratio, defined as the ratio of the inner axonal diameter to the total outer myelinated fiber diameter. Because the g-ratio is a key determinant of neural conduction velocity,^14^ mapping this metric in vivo provides a critical structural mechanism that may in part explain the global functional connectivity disruptions observed in 16p11.2 carriers.^8^

In the present study, we investigate the white matter architecture of 16p11.2 CNV carriers from two independent cohorts across an expanded age range (4 to 60 years). By integrating dMRI and magnetization transfer saturation (MTsat), a marker of myelin density,^15^ we derive estimates for tract axonal volume (AV), myelin volume (MV), and aggregate g-ratio.^16^ This multimodal quantitative MRI framework enables us to evaluate gene-dosage effects and categorical effects across various connectome parcellation, including tract length, canonical resting-state functional networks, and individual cortical nodes. Together, this work provides new insights into how CNVs at the 16p11.2 locus alter white matter microstructure, and more specifically the relationship between axons and myelin.

## Materials and Methods

To investigate the impact of 16p11.2 CNVs on white matter, we analyzed two independent cohorts: the Montreal Genetic Family (MGF) cohort and the Centre Hospitalier Universitaire Vaudois-Lausanne (CHUV-L) cohort. In total, the study included 80 neurotypical controls (38M/42F; age: 36.34 ± 14.32 years; FSIQN=74: 100.79 ± 13.55), 26 deletion carriers (11M/15F; age: 19.89 ± 13.07 years; FSIQN=17: 72.37 ± 17.89), and 13 duplication carriers (9M/4F; age: 25.78 ± 17.90 years; FSIQN=13: 75.63 ± 19.31). Site-specific demographic breakdowns are summarized in Supplementary Table S1. Primary analyses were performed on ComBat-harmonized data from both sites.^17^ To ensure the findings were not driven by site-specific effects, all microstructural analyses were also performed separately for each cohort and are presented in the Supplementary Materials.

### MRI Data Acquisition

The study included two independent cohorts: the MGF and the CHUV-L datasets, comprising individuals who underwent genetic testing. The study was approved by the Research Ethics Board of McGill University (Canada). Data acquisition were approved by local institutional ethics review board (CHU Sainte-Justine and University of Lausanne), and written informed consent was obtained from all adult participants or legal guardians of minors.

Imaging was performed on 3T Siemens Prisma scanners with 64-channel head coils at the McConnell Brain Imaging Centre (Montreal Neurological Institute; MGF) and the Centre Hospitalier Universitaire Vaudois (CHUV-L). Both sites acquired comparable protocols including multi-shell DWI and quantitative MTsat (Table S2-S3).

### MRI Data Processing

Due to differences in acquisition protocols between sites, structural and diffusion preprocessing pipelines were adapted separately for each dataset.

#### Structural Preprocessing

For the MGF dataset, T1w and T2w images were processed with FreeSurfer v7.0’s^18^ standard *recon-all* pipeline to obtain cortical surfaces and parcellations.

For the CHUV-L dataset, which lacked a conventional T1w MPRAGE, the MTsat image (combined with the third echo of the MToff acquisition to improve contrast) was used as the anatomical input. Since FreeSurfer’s *recon-all* performed poorly on these contrast properties, cortical surface reconstruction and parcellation were run with FastSurfer,^19,20^ where the results were visually inspected for quality.

For both datasets, subcortical segmentation were derived using FSL FIRST (v6.0.3),^21^ and tissue-type classification was performed with FSL FAST.^22^ A five-tissue-type segmentation image was then generated for anatomically constrained tractography.^23^

#### Diffusion Preprocessing

Preprocessing was performed in native DWI space using MRtrix3 (v3.0.3).^24^ The pipeline included: image denoising,^25–27^ Gibbs ringing correction (performed using the method of Lee *et al.*^28^ for MGF, and *mrdeggibs*^29^ for CHUV-L to account for the different Partial Fourier factors), susceptibility, motion, and eddy-current correction using FSL TOPUP and EDDY.^30–32^ For the MGF dataset, seven reverse-phase-encoded b=0 images were used for distortion correction. For the CHUV-L dataset, which lacked such pairs, a synthetic distortion-free b0 was generated^33^ to enable TOPUP prior to EDDY. Finally, B1 bias-field correction^34^ was applied. The preprocessed DWI data were upsampled to match the resolution of their respective anatomical images (0.9 mm for MGF and 1.5 mm for CHUV-L). Multi-shell, multi-tissue response functions were then estimated for constrained spherical deconvolution.^35,36^ followed by intensity normalization across tissues.^37,38^ The respective anatomical images (T1w for MGF and MTsat for CHUV-L) were nonlinearly registered to DWI space using ANTs.^39^ Across cohorts, MTsat maps were computed following the Helms model^15^ with B1⁺ correction, implemented via qMRLab^40^ for MGF and via hMRI toolbox^41,42^ for CHUV-L.

### Quantitative Map Computation

Intra-axonal volume fraction (IAVF) maps were obtained by fitting NODDI^43^ using the AMICO toolbox^44^ to the multi-shell diffusion data. MTsat maps were converted into myelin volume fraction (MVF) maps following independent calibration for each site due to differences in the MT acquisition parameters, assuming a reference g-ratio of 0.7 in the healthy young adult splenium^45^. Using the MVF maps together with IAVF, the g-ratio and the axonal volume fraction (AVF) maps were computed.

### Whole Brain Global Models

Whole-brain volume was quantified by summing the white matter, subcortical gray matter and cortical gray matter partial volume estimates from the five-tissue-type segmentation, scaled by voxel dimensions. To derive global microstructural measures free from partial volume effects, analysis was restricted to a conservative ’pure’ white matter mask (voxels with 100% white matter probability). The mean values for AVF, MVF, and g-ratio were computed specifically within this mask and harmonized across sites using neuroCombat.^17^

For each volumetric or microstructural metric (*Y*), a normative baseline model was established using the control cohort from both sites (*Y*∼*Age* + *Age*^2^ + *Sex*). Using the coefficients derived from this healthy baseline, we computed partial residuals (*Y_adj_*) for the entire dataset by regressing out the confounding effects of age (linear and quadratic), while retaining the variance associated with sex. These partial residuals were subsequently analyzed using two distinct linear models (*Y_adj_* = *Sex* + *CNV* + *Sex* × *CNV*) to evaluate the impact of CNV status, biological sex, and their interaction. First, a categorical model was employed to assess whether deletion or duplication carriers differed significantly from the control group. Second, to test for a linear gene-dosage effect, we fitted a continuous model using the same formula but with CNV status encoded as an ordinal variable where deletions were assigned -1, controls were assigned 0, and duplications were assigned +1. This framework allowed us to determine not only if CNVs altered microstructure but also if these effects were sexually dimorphic.

### Tract-Specific Microstructure

Anatomically constrained tractography was performed on the normalized white matter fiber orientation distributions image using the probabilistic algorithm iFOD2.^23,46^ A tractogram of 3 million streamlines was generated using the following parameters: min/max length = 10/400, max angle = 22.5, step = 0.5, cutoff = 0.06, backtrack, crop_at_gmwmi (gray-matter-white-matter interface). The brain was parcellated into 214 nodes using the Schaefer-200 atlas^47^ combined with seven bilateral subcortical regions (amygdala, thalamus, caudate, nucleus accumbens, putamen, hippocampus, pallidum). To mitigate the noise present in individual streamlines, we defined a tract as a cluster of streamlines sharing similar trajectories and/or shared cortical terminations. For each tract, the AV, MV and g-ratio were computed^16^ with the use of the COMMIT framework.^44^

These structural connectomes underwent a filtering and harmonization pipeline:

1. Retain edges present in at least 50% of controls in both MGF and CHUV-L datasets.

2. Cross-site harmonization using ComBat,^17^ applied independently to connectivity matrices for AV, MV, axonal cross-sectional areas, myelin cross-sectional areas, aggregate g-ratio, and tract length.

3. Normalize total edge caliber (summed axonal and myelin cross-sectional areas) at the subject level and retain the top 25% of connections pooled across the cohort. Then, apply a global consensus mask requiring edge presence in at least 50% of subjects to generate the final connectivity matrices.

### Multi-Scale Connectome Modelling

#### Connectome Partitioning

To evaluate microstructural features across spatial and organizational scales, connectivity matrices were partitioned into three analytical frameworks. First, to assess length-dependent effects, harmonized tract length was modeled in controls using a linear mixed-effects model (fixed effects: age, sex; random effects: subject, edge) to derive a standardized edge rank.

Edges were ordered along this spectrum and segmented into bins using a sliding-window approach (window size: 10th percentile; step size: 5th percentile). Second, for resting-state functional network-level analysis, tracts were grouped into an eight-module parcellation consisting of a single unified subcortical module (pooling all subcortical regions) and the seven canonical cortical resting-state networks, evaluating all intra- and inter-network connections. Third, for nodal-level analysis, tracts were aggregated based on their structural connectivity to each specific node, pooling all pathways connected to a given brain region. Across all partitions, we computed total AV (sum), total MV (sum), and cross-sectional-area-weighted mean g-ratio.

#### Normative Modeling and Statistical Analysis

All microstructural metrics were evaluated using a normative modeling. Target variables were first z-scored relative to the control group, and normative baseline were fit on the control cohort to regress out confounding covariates. For volumetric measures (AV and MV), the normative model adjusted for age, sex, and total brain volume (*Y*∼*Age* + *Age*^2^ + *Sex* + *BrainVolume*). For the g-ratio, sex was excluded based on previous literature demonstrating a lack of sexual dimorphism,^48^ and total brain volume was excluded because the g-ratio represents a spatially invariant proportion (*Y*∼*Age* + *Age*^2^). The coefficients derived from this healthy baseline were then applied to the entire cohort to yield covariate-adjusted partial residuals. These standardized residuals were evaluated using general linear models without interaction terms under two frameworks: a categorical model (*Y_adj_*∼*CNV_Categorical_*) to assess independent group differences against the control baseline, and a continuous model (*Y_adj_*∼*CNV_GeneDosage_*) to test for the linear gene-dosage effect. Statistical significance across all length windows, networks, and nodes were determined after correcting for multiple comparisons using the False Discovery Rate (FDR) method.

### Analysis of Other CNVs

To test for shared microstructural alterations across diverse genetic variants, we analyzed a heterogeneous cohort of 29 individuals carrying 25 distinct neurodevelopmental CNVs from the MGF dataset (14M/15F; age: 25.08 ± 14.78 years; FSIQN=28: 81.81 ± 17.74; detailed clinical and genetic characteristics in Supplementary Table S4). Because these variants may exert microstructural effects in different directions, conventional linear models are poorly suited to detect a shared association. We therefore used the Optimal Sequence Kernel

Association Test (SKAT-O), which tests whether CNV carriers exhibit greater phenotypic variability than controls without assuming a common direction of effect. Covariate adjustments and normative baseline modeling remained identical to the primary 16p11.2 pipeline. To ensure that significant variance associations were not driven by single individuals or specific high-impact variants, all analyses were coupled with a leave-one-out (LOO) sensitivity analysis.

## Results

### Negative 16p11.2 gene dosage effect on white matter volume

We first evaluated macroscopic differences in 16p11.2. Total brain (TB), cortical gray matter (cGM), subcortical gray matter (sGM), and white matter (WM) volumes were assessed against a control-derived normative model (Figure S1) incorporating age, age^2^, and sex (Figure 1A). Total brain volume exhibited a significant inverse gene-dosage effect (*p_gene_*_.*TB*_ = 0.0215), characterized by a trend toward larger volumes in deletion carriers (*p_del_*_.*TB*_ = 0.186) and significantly smaller volumes in duplication carriers (*p_dup_*_.*TB*_ = 0.0489). While cGM showed no significant dosage or group differences and sGM reduction were restricted to duplication carriers (*p_dup_*_.*sGM*_ = 0.0153). WM volume exhibited a significant inverse gene-dosage effect (*p_gene_*_.*WM*_ = 0.00245). The categorical results confirmed this effect is symmetrically driven by volume increase in deletion carriers (*p_del_*_.*WM*_ = 0.0448) and reduction in duplication carriers (*p_dup_*_.*WM*_ = 0.0236). These global anatomical findings were consistent across both cohorts (Supplementary Figs. S3A–S4A).

**Figure 1:**
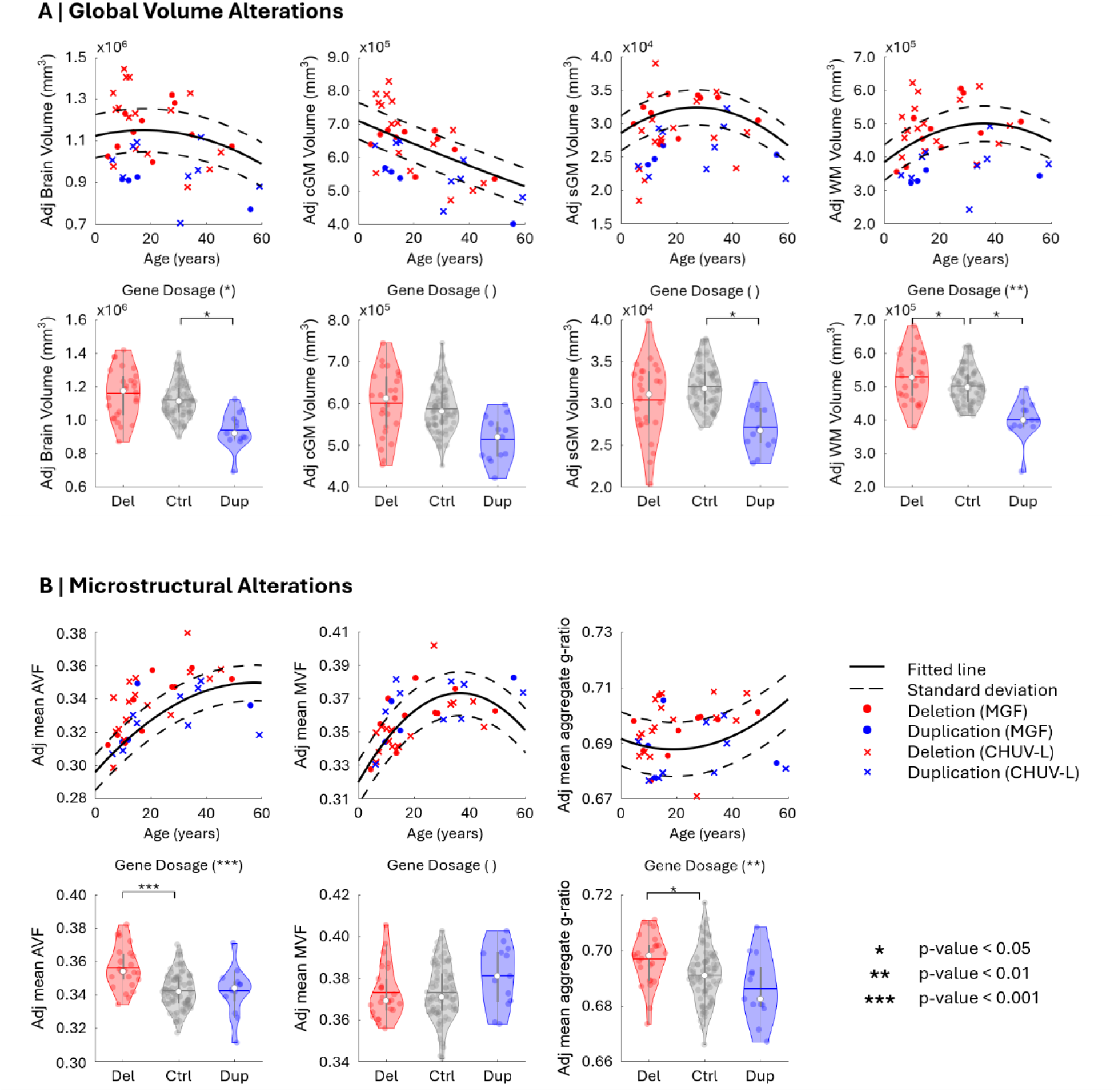
Global macro-anatomical and white matter microstructural alterations across the 16p11.2 CNV spectrum. **(A) Standardized deviations (z-scored) for global volumes (total brain volume, cortical gray matter, subcortical gray matter, and white matter) relative to a normative control baseline.** Analysis revealed significant gene-dosage effects for total brain volume and white matter volume. Specifically, duplication carriers exhibited significant effects in subcortical gray matter and white matter, whereas deletion carriers showed significant effects exclusively in white matter. **(B) White matter microstructural alterations.** Continuous gene-dosage models revealed significant linear relationships for axonal volume fraction and g-ratio, with deletion carriers specifically exhibiting significant effects across these metrics. Conversely, all other comparisons, including all duplication pairwise tests and all models for mean myelin volume fraction, remained non-significant. Notably, a significant Sex × CNV interaction was observed exclusively for the axonal volume fraction gene-dosage model.

### 16p11.2 CNVs disrupt global axonal density and g-ratio, but myelin density is preserved

We next asked whether the reciprocal changes in WM volume reflected alterations in the axonal compartment, the myelin compartment, or their structural relationship. To address this, we quantified the AVF, MVF, and the aggregate g-ratio across all white matter voxels. AVF exhibited a significant inverse gene-dosage effect (*p_gene_*_.*AVF*_ < 10^−4^), driven by higher axon density in deletion carriers (*p_del_*_.*AVF*_ < 10^−4^). The only significant sex × copy number interaction observed across all macroscopic and microstructural metrics was for AVF, specifically within the gene-dosage model (*p* = 0.05). However, the limited sample size precluded reliable sex-stratified analyses to further characterize this effect.

In contrast, MVF showed no significant gene-dosage or categorical differences from the normative baseline. The aggregate g-ratio exhibited a significant inverse gene-dosage effect (*p_gene_*_.*gratio*_ < 0.01), driven primarily by deletion carriers (*p_del_*_.*gratio*_ = 0.0400, *p_dup_*_.*gratio*_ = 0.168). These results suggest that although total white matter MV remains largely stable, the relative myelin sheath thickness is altered in 16p11.2 CNV carriers. These global microstructural findings were broadly preserved across both cohorts (Supplementary Figs. S3B–S4B).

### 16p11.2 gene dosage drives a pervasive, length-independent disruption of the g-ratio

We next asked whether the observed microstructural phenotype preferentially affected short range connections (e.g., U-fibers) or long-range association or interhemispheric pathways. AV exhibited a significant inverse gene-dosage effect that was most pronounced in shorter tracts (Figure 2). Deletion carriers exhibited significantly greater AV within shorter tracts, whereas duplication carriers demonstrated lower AV primarily within medium-and long-range tracts. In contrast, MV remained stable across tract lengths, with no significant gene-dosage or categorical effects. The aggregate g-ratio exhibited a robust inverse gene-dosage effect across all tract lengths. This pervasive, length-independent shift was mirrored in both CNV groups: deletion carriers demonstrated significantly higher g-ratio (thinner relative myelin sheath), while duplication carriers exhibited significantly lower g-ratios across nearly all bins (thicker relative myelin sheath). These tract length-dependent profiles were consistent across both cohorts (Supplementary Figs. S5–S6).

**Figure 2:**
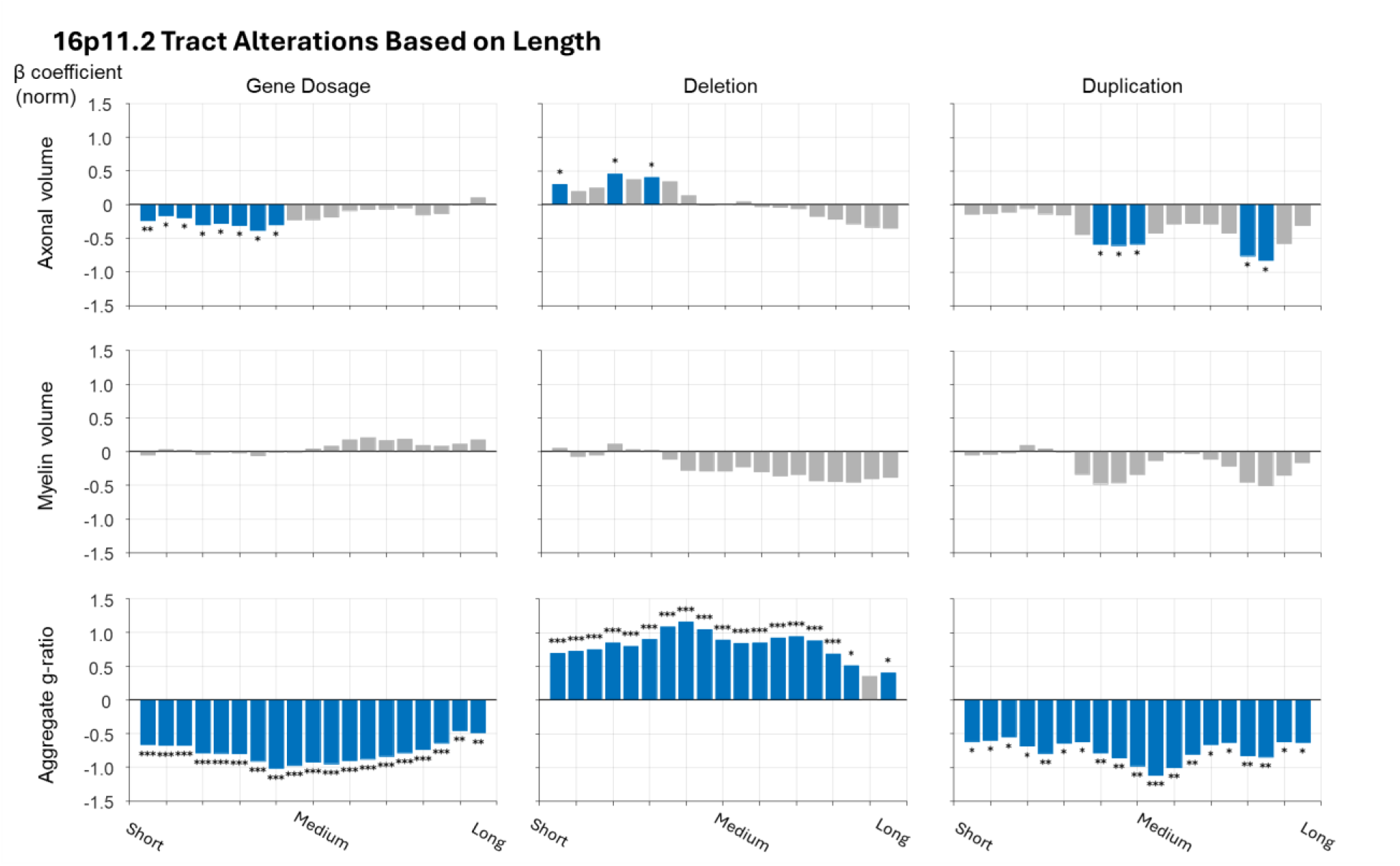
Distribution of gene-dosage effect and categorical effect sizes across the tract length spectrum. Standardized effect sizes for **axonal volume** (top row), **myelin volume** (middle row), and **aggregate g-ratio** (bottom row). The x-axis represents tracts ranked by length and grouped into moving-window bins of 10%. Bar heights represent linear model effect sizes for each bin; gray bars indicate results that did not survive FDR correction. Notably, while the magnitude of effect sizes for both axonal volume and the g-ratio are comparable between deletion and duplication carriers, the statistical significance of the g-ratio alterations is substantially higher in the deletion cohort.

### Deletion carriers drive widespread gene-dosage effects on the aggregate g-ratio

We next asked how these microstructural alterations map onto the brain’s functional architecture by evaluating connections within and between canonical resting-state networks (Figure 3). Network-level analysis revealed a significant inverse gene-dosage effect for AV, most prominently within the Visual, Salience/Ventral Attention, and Limbic networks. Deletion carriers exhibited widespread increases in AV (although a small subset of connections showed lower AV), whereas duplication carriers exhibited lower AVs that were localized to subcortical connections. MV exhibited minimal deviation from the normative baseline, with substantially fewer significant differences than AV. All significant effects identified in the continuous gene-dosage model were also observed in deletion carriers, whereas duplication carriers showed a significant MV difference only within the subcortical network.

**Figure 3:**
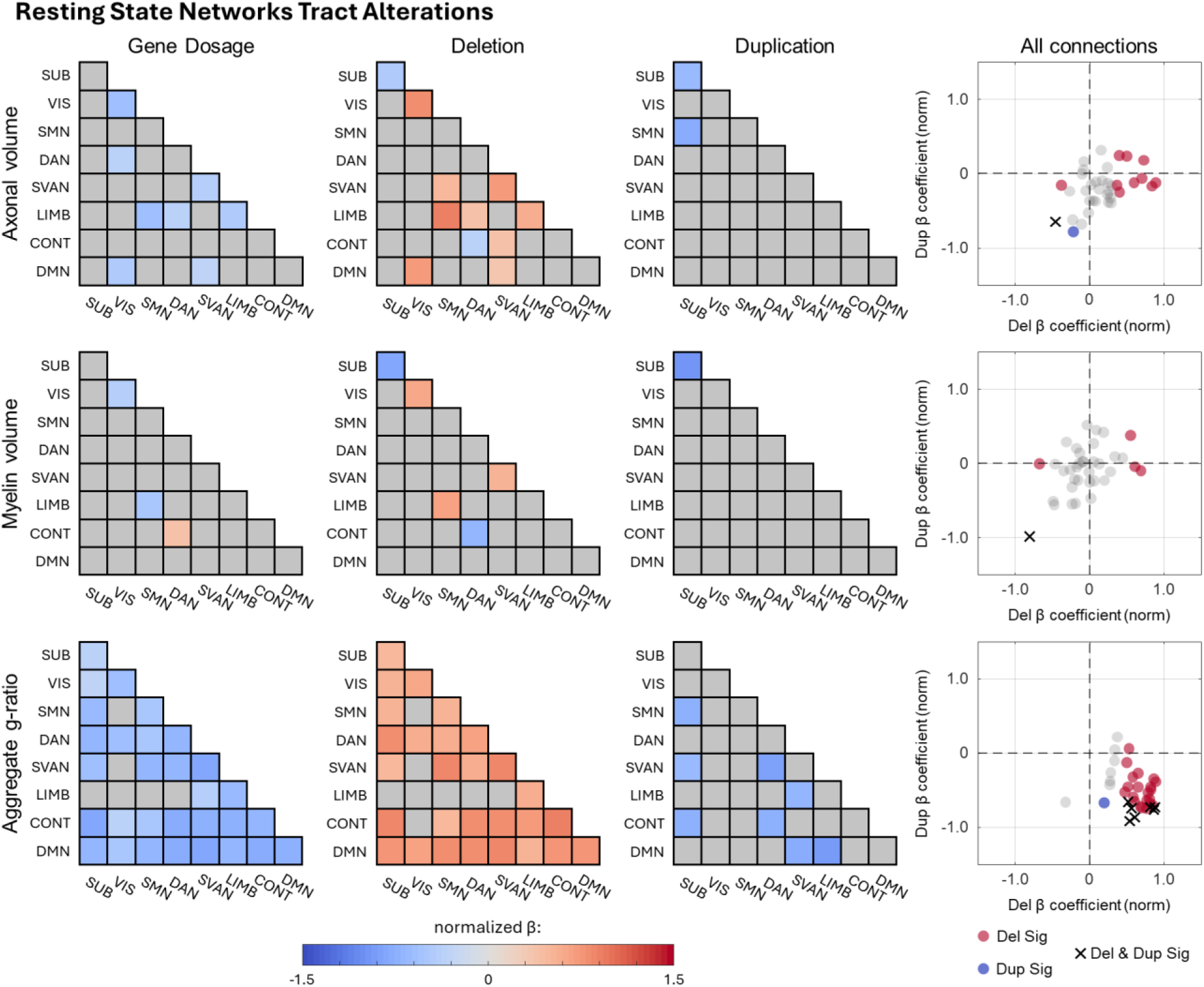
Distribution of gene-dosage effects and categorical effects across functional networks. Significant gene-dosage effects were predominantly observed for both axonal volume and the aggregate g-ratio, while myelin volume exhibited minimal deviation. The color scale represents standardized z-scores relative to the normative baseline, where gray regions indicate results that did not reach significance after FDR correction. The rightmost column summarizes categorical significance: red indicates regions significantly altered in deletion carriers, blue indicates significance in duplication carriers, purple crosses denote significance across both groups, and gray denotes no significant categorical effect.

The aggregate g-ratio exhibited a strong, network-wide inverse gene-dosage effect, except for some limbic and visual connections. The deletion cohort yielded significant positive g-ratio deviations between and within most functional networks. Although fewer connections reached significance in duplication carriers, the direction of the effect consistently mirrored the reciprocal gene-dosage pattern, manifesting as lower g-ratios. These network-level patterns were broadly consistent across both cohorts (Supplementary Figs. S7–S8).

At the node level (Figure 4), continuous modeling revealed a widespread negative gene-dosage effect for AV, predominantly involving the bilateral visual cortex and the inferior frontal gyrus. Deletion carriers exhibited widespread increases in AV together with a smaller subset of nodes showing reduced AV, whereas duplication carriers showed no significant nodal AV alterations. In contrast, MV exhibited high spatial heterogeneity. The continuous gene-dosage model yielded a mixture of localized MV increases and decreases that lacked a cohesive global trajectory. Deletion carriers presented slightly more significant nodes than the continuous model; however, these localized alterations remained directionally inconsistent, and duplication carriers again showed no significant regional MV alterations. The aggregate g-ratio exhibited a widespread inverse gene-dosage effect. Although few nodes reached statistical significance in duplication carriers, the direction of effect consistently aligned with the reciprocal gene-dosage pattern. These profiles were also consistent across both cohorts (Supplementary Figs. S9–S10).

**Figure 4:**
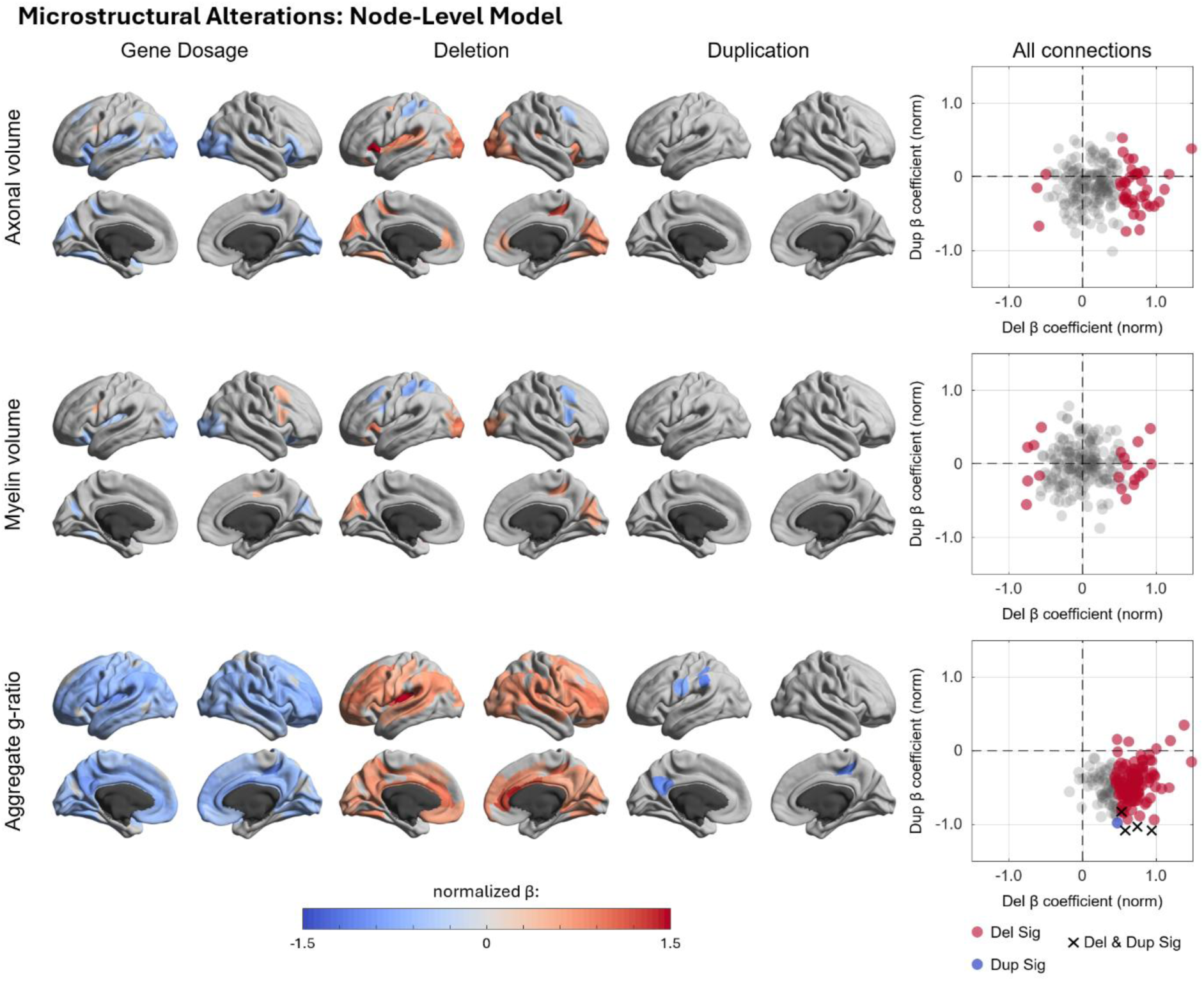
Distribution of gene-dosage effects at nodal scale. Axonal volume (AV) and the aggregate g-ratio exhibited the most widespread alterations in the continuous gene-dosage model, a pattern driven primarily by the deletion cohort. In contrast, alterations in myelin volume were sparse and spatially heterogeneous. The color scale represents standardized z-scores relative to the normative baseline, where gray regions indicate results that did not reach significance after FDR correction. The rightmost column summarizes categorical significance: red indicates regions significantly altered in deletion carriers, blue indicates significance in duplication carriers, purple crosses denote significance across both groups, and gray denotes no significant categorical effect.

Direct comparison of standardized effect sizes via Wilcoxon signed-rank tests confirmed that both AV and g-ratio effect sizes were significantly greater in deletion than duplication carriers (*p_AV_* < 0.001, *p_gratio_* < 0.01). In contrast, MV effect sizes showed bidirectional changes and lacked a systematic gene-dosage pattern.

Collectively across all levels of connectome organization, we observed a highly consistent microstructural phenotype associated with 16p11.2 gene dosage. Despite relatively stable MV, deletion carriers consistently exhibited greater AV alongside a higher aggregate g-ratio, indicating relatively thinner myelin sheaths, whereas duplication carriers exhibited reduced AV and a lower aggregate g-ratio, indicating relatively thicker myelin sheaths.

### 16p11.2 exhibits a similar white matter profile compared to other CNVs

To determine whether the 16p11.2 microstructural phenotype is locus-specific or shared across neurodevelopmental risk variants, we evaluated the heterogeneous CNV cohort using SKAT-O. At the whole brain level (Figure 5A), the heterogeneous CNV cohort exhibited greater variability than neurotypical controls for all tissue volumes. However, LOO analyses indicated that the WM volume was largely driven by a single individual. Among the global WM microstructural metrics (Figure 5A), AVF’s increased variance was also driven by a single individual and rendered non-significant under LOO exclusion, whereas MVF and global aggregate g-ratio showed no significant variance differences.

**Figure 5:**
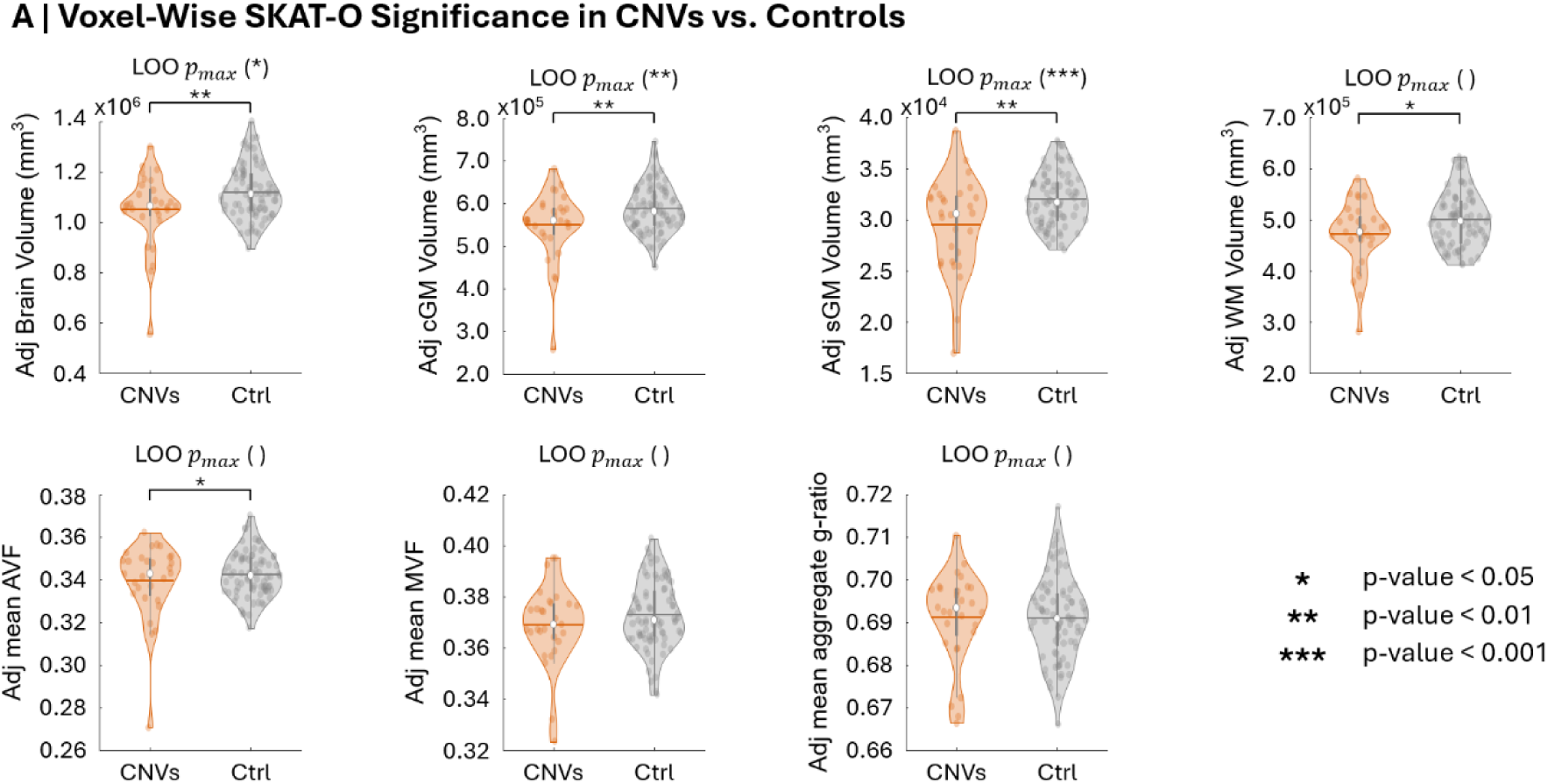
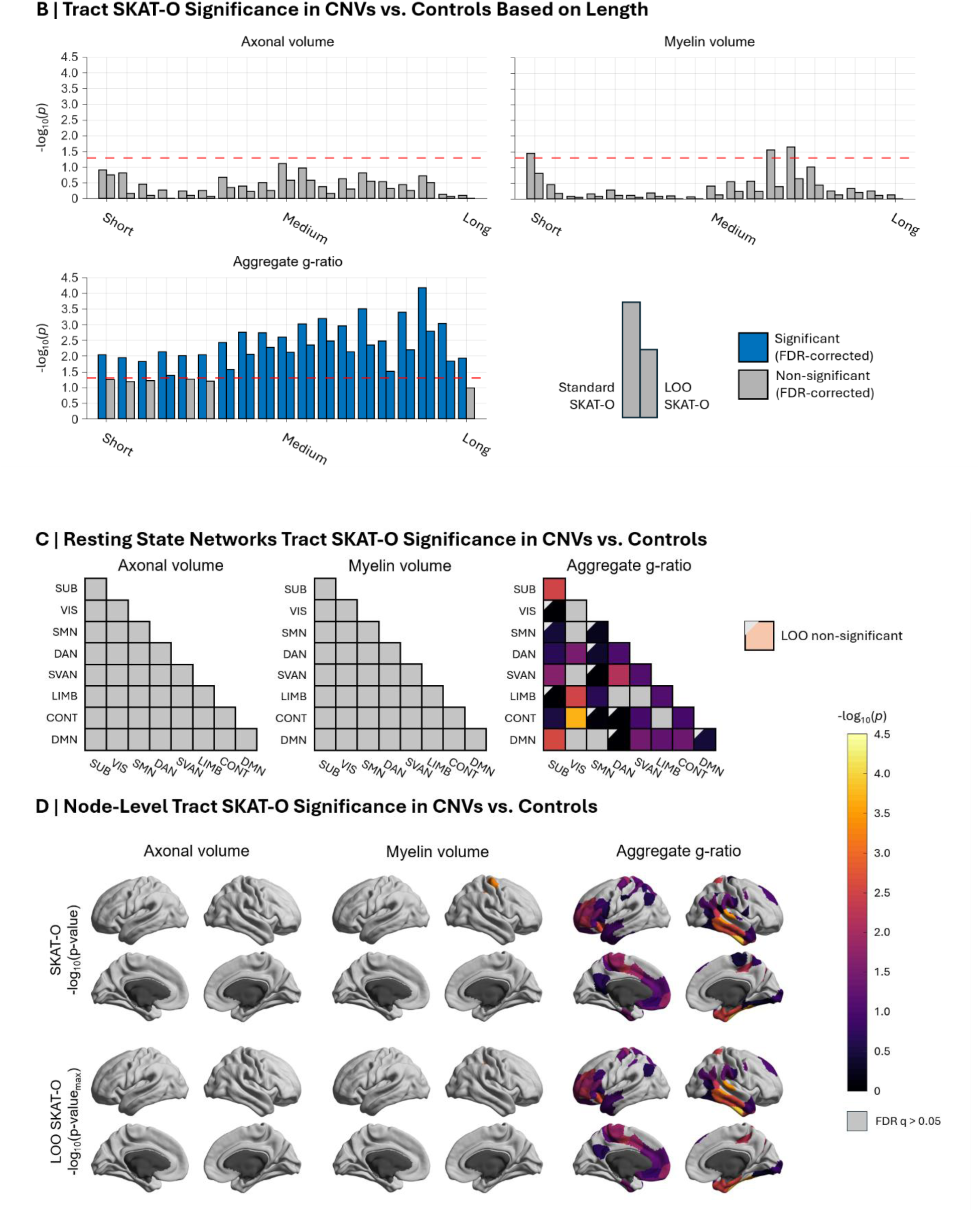
Comparative variance profiling across a heterogeneous cohort of neurodevelopmental CNVs. **(A)** Global SKAT-O variance significance across tissue volumes and whole-brain microstructural metrics, alongside LOO analysis, demonstrating that global alterations are predominantly present in global tissue volumes. **(B)** Tract-length binning of microstructural features, demonstrating variance comparable to neurotypical controls in AV and MV after multiple comparison corrections, contrasting the widespread g-ratio variance alterations. **(C)** Network-level structural variance across canonical network connections; AV and MV variance remain unaltered, whereas altered g-ratio variance connections are predominantly concentrated within higher-order cognitive networks following LOO validation. **(D)** Cortical node-level mapping, illustrating an absence of variance effects in AV, isolated alterations in MV, and widespread g-ratio variance differences localized within higher-order association cortices while largely excluding primary sensory and motor regions.

Across tract lengths (Figure 5B), AV and MV showed no significant differences in variability across lengths relative to controls. In contrast, the aggregate g-ratio exhibited significantly greater variance across nearly all tract-length bins, remaining robust under LOO sensitivity (becoming marginally non-significant only within the shorter and the longest bin).

Topological analyses at both the network and node level closely mirrored this pattern. While AV and MV variability remained comparable to controls across both scales, aggregate g-ratio variance was widely elevated. However, following LOO analysis, these effects became concentrated within higher-order association networks, specifically the salience/ventral attention, limbic, and control networks.

## Discussion

Our findings demonstrate that 16p11.2 CNVs are associated with a pervasive, gene-dosage dependent white matter microstructural phenotype. Reciprocal alterations in global white matter volume were consistent with previous large-scale studies.^6,49^ Although total brain and cortical gray matter volumes did not reach statistical significance, both exhibited the same reciprocal trend reported previously, with deletions trending toward higher volumes and duplications toward lower volumes. This discrepancy may be driven by age differences: prior work evaluated a much younger deletion cohort (under 18 years old) alongside an older duplication cohort. Subcortical volume reductions in duplication carriers strongly agreed with previous work, whereas deletion cohort showed a non-significant trend toward smaller volumes, diverging from the larger volumes previously reported. Overall, although statistical significance differed across tissue compartments, the observed anatomical trends were broadly consistent with previous work.

### Axon-myelin imbalance drives brain-wide g-ratio disruption

Previous diffusion MRI studies consistently reported higher FA in deletion carriers and lower FA in duplication carriers,^9,10^ but FA cannot distinguish whether these differences stem from axons, myelin, and/or fiber geometry. Building on recent evidence of increased intra-axonal volume in young deletion carriers,^12^ we extend these findings into an older cohort while simultaneously examining both deletion and duplication carriers. Across our analyses, we observed a reciprocal gene-dosage effect on tract AV alongside comparatively stable MV. These findings indicate that previously reported diffusion MRI alterations reflect changes in the axonal compartment rather than absolute myelin content. This imbalance manifests as an altered aggregated g-ratio (Figure 6), suggesting a disruption in the normal coordination between axonal growth and myelin remodelling.

**Figure 6:**
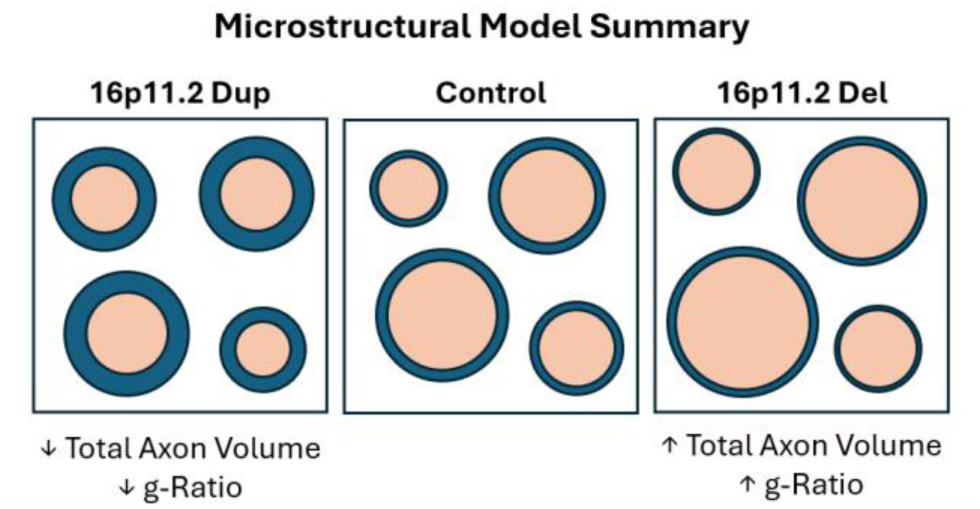
Conceptual model of gene-dosage effects on white matter microstructure. Schematic representation of the reciprocal microstructural alterations associated with 16p11.2 copy number variations. Duplications are characterized by reduced axonal volume and a disproportionately thick myelin sheath relative to axonal diameter. Conversely, deletions are associated with increased axonal volume and a reduced relative myelin thickness. Importantly, because our imaging protocol cannot distinguish between changes in axon count and individual axon caliber, this model assumes the total number of axons remains constant.

The g-ratio is a key determinant of conduction velocity,^14^ reflecting a fundamental trade-off between signal transmission, metabolic efficiency, and spatial constraints.^50^ Disrupted g-ratio has been implicated in a wide range of psychiatric and neurological conditions.^51–53^ As the g-ratio is a relative measure, it is less sensitive to global volumetric differences, making it a robust marker of changes in the relationship between axons and myelin. Because this altered relationship cannot be resolved using diffusion or myelin imaging alone, integrating quantitative measures of both the axonal and myelin compartments was essential to capture the disrupted axon-myelin relationship in the white matter of 16p11.2 CNVs, manifesting as elevated g-ratio in deletions and reduced g-ratio in duplication carriers.

Because our metrics aggregate across whole tracts, this macroscopic profile could arise from several biological scenarios: altered axon number without proportional changes in myelination, shift in average axon caliber, and/or disruption in the distribution of axon calibers within a tract. For example, because larger-caliber axons exhibit higher g-ratios,^54^ selective pruning of large-caliber axons could shift the aggregate g-ratio without uniform changes across the entire axonal population. Distinguishing among these possibilities will be essential for understanding how 16p11.2 gene dosage alter axon-myelin development.

### Increased vulnerability in 16p11.2 deletions

Although deletions and duplications exerted reciprocal effects, categorical analyses revealed substantially more widespread microstructural alterations in deletion carriers, and direct comparisons of nodal effect sizes confirmed that these alterations were significantly greater in magnitude. This pronounced vulnerability to deletions aligns with broader clinical trends, wherein deletion carriers consistently exhibit more severe health and developmental phenotypes than duplication carriers.^2^ This asymmetry is also consistent with animal models, in which 16p11.2 deletion mice exhibit significantly higher rates of premature mortality.^55^ These neuroimaging findings provide structural evidence that deletion carriers are more severely affected than duplication carriers, mirroring the lower average FSIQ observed among deletion carriers in our dataset. This asymmetry is consistent with human inheritance patterns in which deletions arise more frequently de novo whereas duplications are more commonly inherited.^56^

### Shared g-ratio disruption in neurodevelopmental risk

Evaluating a heterogenous CNV cohort revealed that this microstructural phenotype is not unique to the 16p11.2 locus. Absolute AV and MV did not reach statistical significance, this does not rule out underlying differences, which may be obscured by the limited sample size. In contrast, the aggregate g-ratio remained significantly altered, consistent with previous reports of both elevated and reduced g-ratios across diverse neurodevelopmental disorders.^57,58^ Rather than a locus-specific anomaly, altered axon-myelin balance may represent a shared microstructural vulnerability across generic neurodevelopmental risk.

### Robustness across analytical frameworks

Crucially, this microstructural phenotype was robust across both tract-specific and conventional tractometry frameworks (Figure S11). The convergence between these different analytical pipelines indicates that the observed microstructural alterations are not artifacts of a particular modeling assumption but rather reflects a robust biological feature of the 16p11.2 white matter phenotype.

### Limitations

The *in vivo* MRI protocol used in this study provide aggregate measures of tissue microstructure in a whole voxel or tract; it cannot provide a distribution of axon diameters or myelin g-ratios in the voxel or tract. Furthermore, because diffusion-derived metrics are signal-weighted toward large caliber axons,^59^ our aggregate metrics predominantly reflect those dominant fibers rather than the entire axonal population. Disentangling these features *in vivo* will require advanced MRI techniques and biophysical models, including multi-compartment diffusion-T2 relaxometry^60^ and axonal caliber mapping.^61^ Integrating these techniques with early longitudinal developmental data will also be critical as prenatal brain growth is heavily driven by cellular proliferation (which dictates axon count),^62^ whereas later postnatal trajectories are dominated by cellular hypertrophy and myelination (which dictate axon diameter).^63^ Longitudinal neuroimaging studies may help determine when and how 16p11.2 gene-dosage influences white matter development at the cellular level.

Second, while multisite harmonization increased statistical power, datasets combining diffusion and magnetization transfer imaging remain scarce. We observed a significant sex × CNV interaction for whole white matter AVF, but our limited sample size precluded reliable evaluation of sex-specific effects at higher levels of connectome organization. Future large-scale efforts prioritizing high-resolution, multi-modal acquisitions will be crucial to enable robust sex-stratified analyses, which is particularly relevant given the ongoing debate in the literature regarding sex effects in 16p11.2.^64,65^ Furthermore, our cohort is weighted toward older participants; extending this multi-modal framework to younger participants would offer critical insights into how axonal and myelin alterations first emerge and for identifying potential windows for intervention. Finally, higher-resolution imaging may enable a more detailed characterization of cortical microstructure, extending this framework beyond white matter pathways.

Despite these limitations, the consistency of the findings across cohorts and complementary analytical frameworks supports the robustness of the observed 16p11.2 white matter microstructure phenotype. We demonstrate a reciprocal gene-dosage effect in which duplication carriers exhibit reduced tract axonal volumes and a lower aggregate g-ratio, consistent with thicker myelin sheaths relative to axon caliber, whereas deletion carriers exhibit greater tract axonal volume and a higher aggregate myelin g-ratio, consistent with relatively thinner myelin sheaths. Together, these findings identify disrupted axon-myelin relationships as a prominent feature of the 16p11.2 white matter phenotype.

## Data availability

Requests for access to the raw neuroimaging data, or processed datasets used in this study should be directed to the corresponding author. The analysis code can be found in: https://github.com/wendalu/16p11.2-wm-gratio

## Acknowledgements

This work was supported by the following funding sources: Canadian Institute of Health Research (PJT-191838), Fonds de Recherche du Québec—Nature and technologies, Canadian Neurodevelopmental Research Training Platform, Natural Sciences and Engineering Research Council of Canada, Brain Canada, Fonds de Recherche du Québec— Santé, CFREF Healthy Brains Healthy Lives, and the Killam Trusts.

