## Supplemental Material for "Dose-dependent myelin g-ratio alterations in the white matter of 16p11.2 copy number variations"

### Site Specific Demographic Breakdown

**Table S1: Summary of Participant Age and Sex Distribution. FSIQ = Full Scale IQ.**

| | MGF | CHUV-L | Harmonized |
|---|---|---|---|
| Controls | 26M/28F<br>35.73 ± 15.15 years<br>FSIQ: 99.91 ± 12.74 ($N_{IQ}$ = 50) | 12M/14F<br>38.00 ± 12.52 years<br>FSIQ: 106.27 ± 13.55 ($N_{IQ}$ = 24) | 38M/42F<br>36.34 ± 14.32 years<br>FSIQ: 100.79 ± 13.55 |
| 16p11.2 Deletion | 2M/8F<br>21.48 ± 13.75 years<br>FSIQ: 87.55 ± 10.03 ($N_{IQ}$ = 4) | 9M/7F<br>18.91 ± 12.97 years<br>FSIQ: 71.25 ± 16.52 ($N_{IQ}$ = 13) | 11M/15F<br>19.89 ± 13.07 years<br>FSIQ: 72.37 ± 17.89 |
| 16p11.2 Duplication | 4M/0F<br>23.21 ± 21.93 years<br>FSIQ: 76.90 ± 20.74 ($N_{IQ}$ = 4) | 5M/4F<br>26.93 ± 17.18 years<br>FSIQ: 77.56 ± 17.52 ($N_{IQ}$ = 9) | 9M/4F<br>25.78 ± 17.90 years<br>FSIQ: 75.63 ± 19.31 |

### Acquisition Parameters

**Table S2: MRI acquisition parameters for the MGF dataset.**

| **Sequence** | **Resolution ($mm^3$)** | **TR (ms)** | **TE (ms)** | **Flip Angle (°)** | **Acceleration** | **Acquisition Details** | **Acquisition Time (min)** |
|---|---|---|---|---|---|---|---|
| T1w | 0.9 iso | 2300 | 2.32 | 8 | GRAPPA = 4 | TI = 900 ms | 3:26 |
| T2w | 0.9 iso | 3200 | 408 | Variable | GRAPPA = 4 | | 3:10 |
| DWI | 1.7 iso | 4100 | 71 | 90 | GRAPPA = 2<br>MB = 3 | b-values (vols):<br>0 (20), 650 (15), 1000 (30), 2000 (60) s/mm²<br>PF = 6/8 | 11:10 |
| MTsat | 1.7 iso | - | - | | | | 9:54 |
| • MTon | 1.7 iso | 33 | 3.81 | 10 | GRAPPA = 2 | PF = 6/8<br>Offset freq:<br>1200 Hz, 500° | 4:02 |
| • MToff | 1.7 iso | 33 | 3.81 | 10 | GRAPPA = 2 | PF = 6/8 | 4:02 |
| • T1w | 1.7 iso | 15 | 3.81 | 25 | GRAPPA = 2 | PF = 6/8 | 1:50 |
| $B1^+$ map | 2 x 2 x 5 | 4000 | 18 | 60/120 | | | 2:16 |

**Table S3: MRI acquisition parameters for the CHUV-L dataset.**

| Sequence | Resolution ($mm^3$) | TR (ms) | TE (ms) | Flip Angle (°) | Acceleration | Key Acquisition Details | Acquisition Time (min) |
|---|---|---|---|---|---|---|---|
| DWI | 2 iso | 7400 | 69 | 90 | GRAPPA = 2 | b-values (vols): 0 (13), 650 (15), 1000 (30), 2000 (60) s/mm² | 14:55 |
| MTsat | 1.5 iso | - | - | | | | 9:21 |
| • MTon | 1.5 iso | 24.5 | 2.34 to 14.04 | 6 | GRAPPA = 2 | PF = 6/8<br>Offset freq:<br>1200 Hz, 500° | 3:07 |
| • MToff | 1.5 iso | 24.5 | 2.34 to 18.72 | 6 | GRAPPA = 2 | PF = 6/8 | 3:07 |
| • T1w | 1.5 iso | 24.5 | 2.34 to 18.72 | 21 | GRAPPA = 2 | PF = 6/8 | 3:07 |
| $B1^+$ map | 4 iso | - | - | | | | 2:45 |
| • B1 ref | 4 iso | 200 | 19.53 /39.06 | Variable | GRAPPA = 2 | | 1:12 |
| • GRE | 3 x 3 x 2.5 | 700 | 4.92/ 7.38 | 80 | | | 1:33 |

### MGF Dataset CNV Subset

**Table S4: Detailed composition of the heterogeneous CNV comparison cohort.**

| CNV Locus | N (Subjects) |
|---|---|
| Del14q32.32-33 | 1F |
| Del18p terminale | 1F |
| Del19p13.3 | 1F |
| Del1q21.1 | 1M/2F |
| Del22q11.2 | 1F |
| Del6p25.1p24.2 | 1M |
| Del7q35 | 1M |
| DelXp22.11 | 1F |
| Dup12p13.31 | 1M |
| Dup14q11.2 | 1M |
| Dup15q11.2, Del 1q21.1 | 1F |
| Dup15q11.2q13.1 | 1M |
| Dup15q11.3q13.1 | 1M |
| Dup15q25.2q25.3 | 1F |
| Dup16p13.11 | 1M/1F |
| Dup17q12 | 1M |
| Dup1q21.1 | 1F |
| Dup22q11.2 | 1M |
| Dup22q11.21 | 1F |
| Dup22q13.31 | 1M |
| Dup2q11.2 | 1M |
| Dup8p21.2p21.1 | 1F |
| Dupxq25 | 1F |
| DupXq28 | 1M/1F |
| SNV (CHD3) | 1M |

### Validation of Normative Model

This section visualizes the normative models (Figure S1) applied to the harmonized control cohort, demonstrating that the expected age-related microstructural effects are properly captured to establish a reliable baseline for our primary analyses.

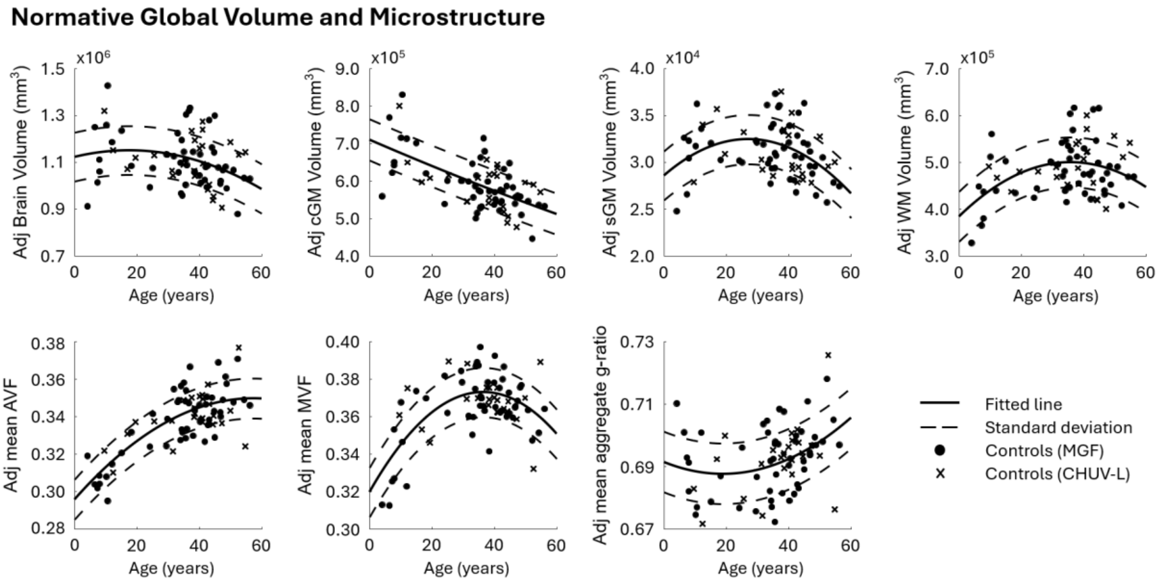


***Figure S1: The fitted normative trajectories are plotted for the combined control cohort.*** *Data were adjusted for site-specific scanner effects utilizing location and scale adjustments via neuroCombat. Normative models, incorporating both ag,* $age^2$*, and biological sex as covariates, were then fitted to establish baseline distributions across all seven metrics evaluated in the main manuscript (e.g., total brain volume, cortical gray matter volume, subcortical gray matter volume, white matter volume, mean axonal volume fraction, mean myelin volume fraction, and mean aggregate g-ratio). The main effects of sex were regressed out to isolate and plot the underlying age-related trajectories. These control-exclusive models serve as the reference distributions for calculating z-scores for individuals with CNVs in the main analyses.*

### Methodological Validation and Consistency with Established IAVF Frameworks

Previous research in 16p11.2 has primarily utilized the Intra-axonal Volume Fraction (IAVF).[1] While IAVF captures essential microstructural trends, it is inherently limited by the insensitivity of standard diffusion MRI sequences to the myelin signal. Because the volume occupied by the myelin sheath is not explicitly modeled, IAVF estimates are often systematically inflated. In this work, we transition to the Axonal Volume Fraction (AVF), which incorporates the Myelin Volume Fraction (MVF) to provide a more biologically specific estimate of axonal space.

To ensure our findings remain grounded in the existing literature, we compared both metrics (Figure S2). The IAVF data demonstrate general trends consistent with previous studies; specifically, deletions are associated with an increase in IAVF. This directional effect, alongside significance levels and effect sizes, is similar to our AVF analysis. However, a notable divergence between the metrics emerges in older age cohorts. This subtle shift is expected, as AVF explicitly captures the dynamic changes in myelination that occur during later developmental and aging stages, which IAVF conflates. Ultimately, the strong baseline correspondence between these models serves as mutual validation, confirming that our results are consistent with established 16p11.2 findings while offering improved specificity regarding age-dependent myelin trajectories.

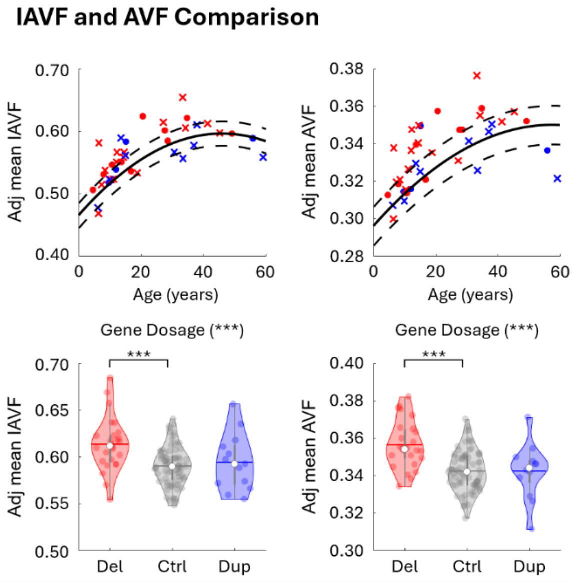


***Figure S2: Linear model comparison of IAVF (left) and AVF (right).*** *Both models demonstrate convergent statistical profiles and directional trends, with a significant increase in axonal proxies observed in 16p11.2 deletion carriers. The strong correspondence in p-values and effect sizes between the two models confirms that our results are robust and fundamentally consistent with prior research, validating that the transition to a more physically specific AVF framework maintains continuity with established biological trends.*

### Validation of Core Findings Across Independent Cohorts

Overall, the site-specific analyses (Figure S3-S10) demonstrate general agreements with the primary harmonized results. As anticipated, we observed minor deviations between the independent cohorts, primarily localized to absolute axonal and myelin volume metrics. This variability is expected as absolute volumetric measures are inherently spatially variant. Within smaller, isolated cohorts, natural inter-subject differences in baseline regional parcellation sizes make absolute volumes highly susceptible to morphological noise, which can cause estimated effect sizes to naturally fluctuate.

In contrast, the g-ratio is a microstructural proportion rather than an absolute volume. This acts as a spatially invariant measure, rendering it significantly more robust against regional morphological differences. As a result, the g-ratio results demonstrated strong agreement across both independent datasets. Specifically, deletions exerted a significant positive effect on the g-ratio, and duplications exhibited a negative, albeit non-significant, effect. Additionally, these site-specific analyses also replicated the significant inverse gene dosage effect observed in the main manuscript, confirming that our findings are stable across both datasets.

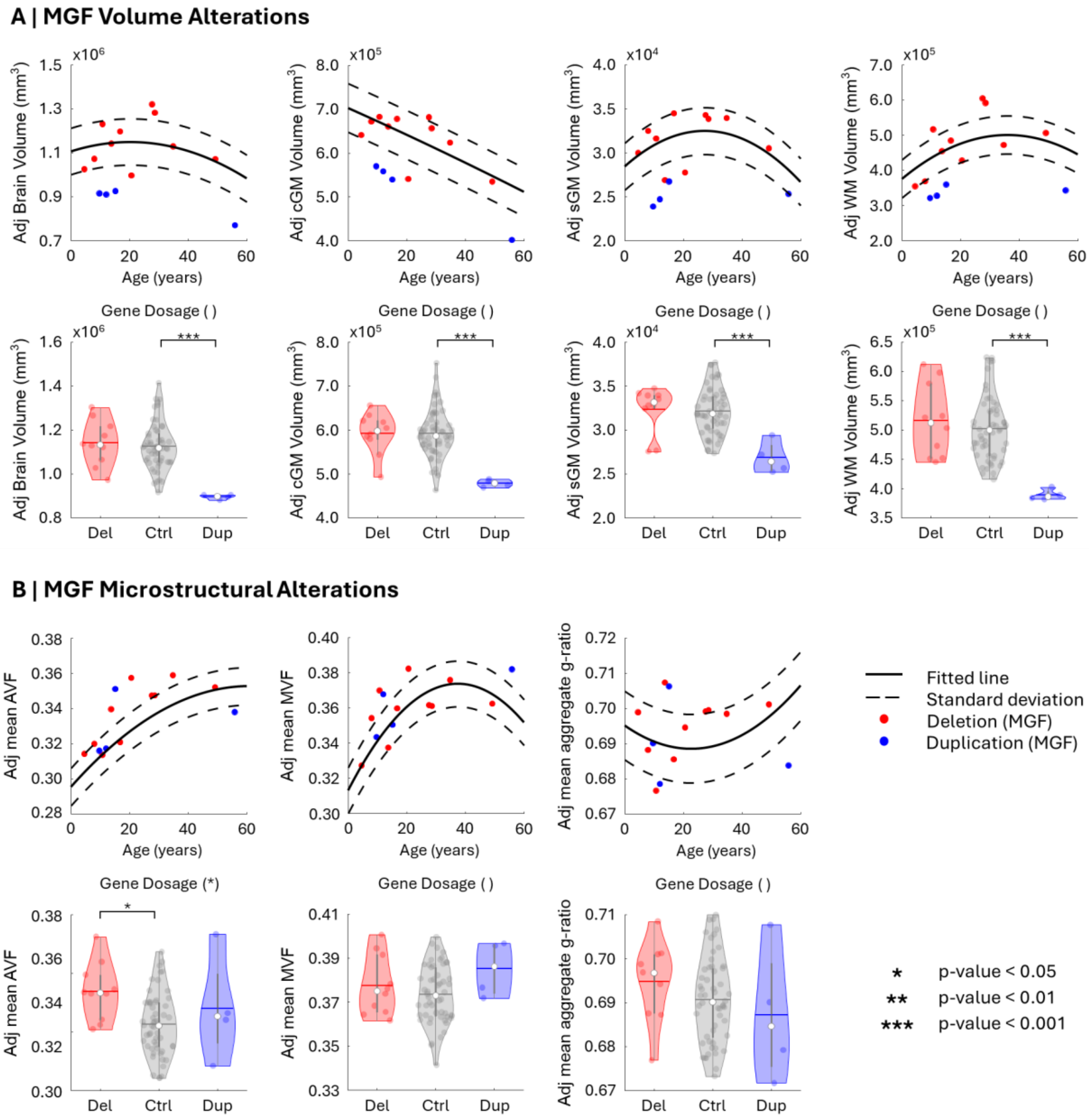

**Figure S3: Global macro-anatomical and white matter microstructural alterations across the 16p11.2 CNV spectrum for the MGF dataset. (A) Standardized deviations (z-scored) for global volumes (total brain volume, cortical gray matter, subcortical gray matter, and white matter) relative to its normative control baseline.** Analysis revealed a significant duplication effect across all volumetric measures, while no significant effects were observed for deletions or the continuous gene-dosage models. **(B) White matter microstructural alterations.** Gene-dosage and categorical deletion models revealed significant relationships only for the mean AVF. Conversely, all other microstructural comparisons, including all pairwise duplication tests and all models for mean MVF and aggregate g-ratio, remained non-significant. Notably, while the mean values for the aggregate g-ratio follow the expected gene-dosage trend described in the main manuscript, these effects did not reach statistical significance in this dataset. This lack of significance is likely attributable to high intra-group variance within the 16p11.2 cohorts at this site, which limited the statistical power to resolve these microstructural deviations.

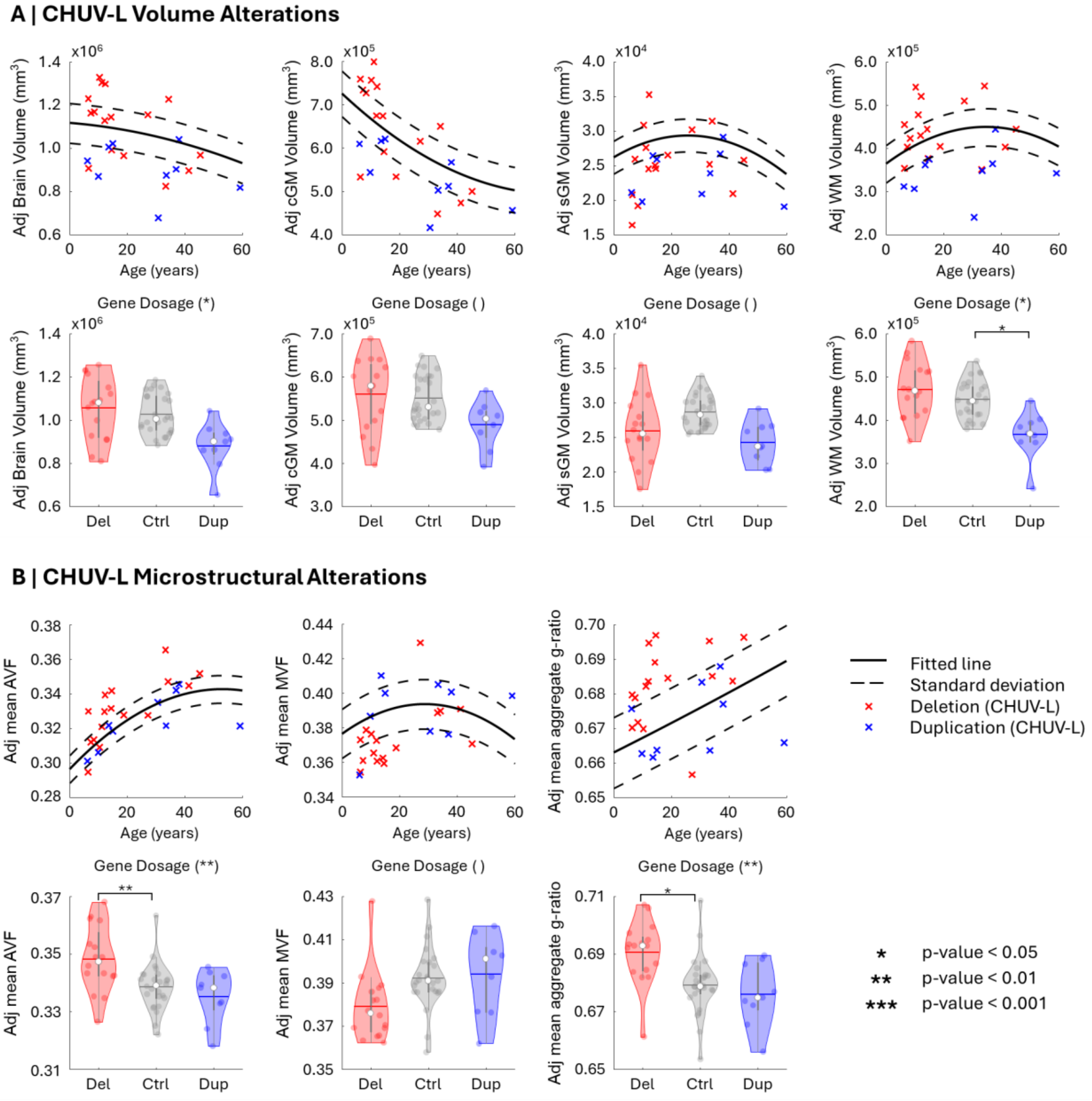


**Figure S4: Global macro-anatomical and white matter microstructural alterations across the 16p11.2 CNV spectrum for the CHUV-L dataset. (A) Standardized deviations (z-scored) for global volumes (total brain volume, cortical gray matter, subcortical gray matter, and white matter) relative to its normative control baseline.** Analysis revealed a significant gene dosage effect for total brain volume and white matter volume, alongside a significant categorical difference between duplication carriers and controls for white matter volume. **(B) White matter microstructural alterations.** Continuous gene-dosage and categorical deletion models revealed significant relationships for both the mean axonal volume fraction and the aggregate g-ratio. Conversely, all other microstructural comparisons, including all pairwise duplication tests and all models for mean myelin volume fraction, remained non-significant. Notably, these results demonstrate strong agreements with the findings reported in the main manuscript.

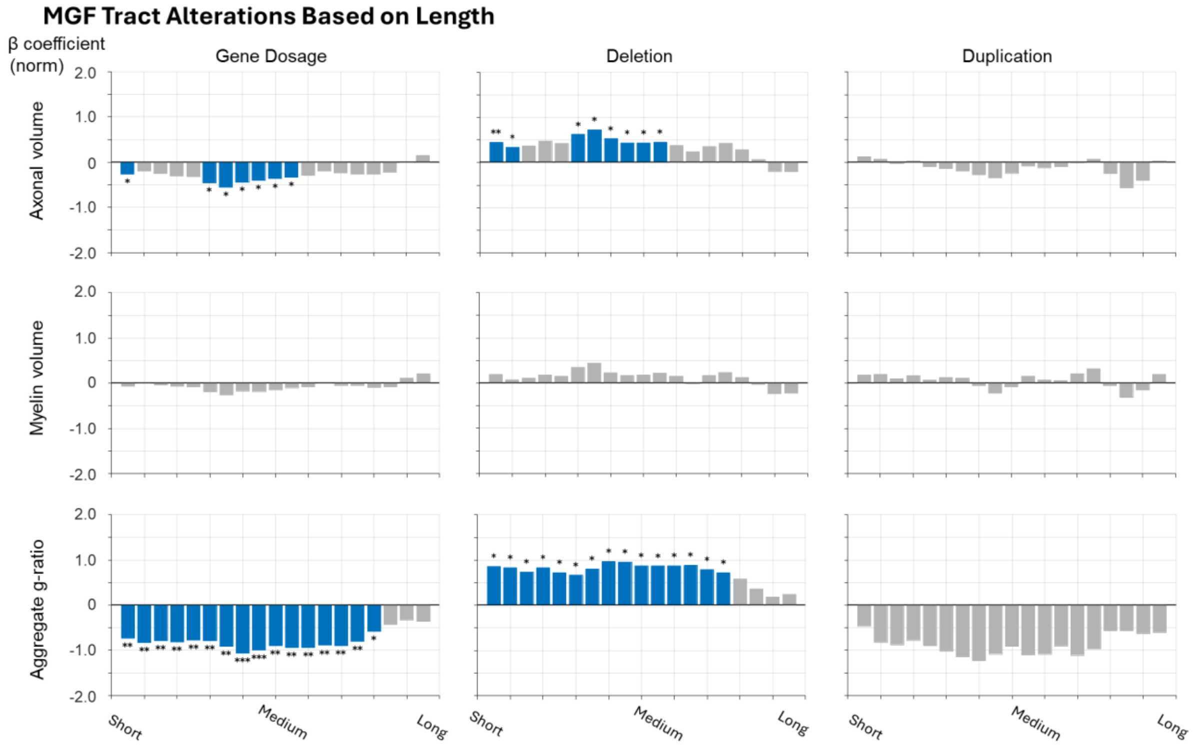


**Figure S5: Distribution of microstructural gene-dosage and categorical effect sizes across the tract-length spectrum in the MGF dataset.** The x-axis represents white matter tracts ranked by physical length and grouped into 10% sliding-window bins. Bar heights represent standardized effect sizes (z-scores) for the continuous gene-dosage model and categorical groups; gray bars indicate results that did not survive FDR correction. The results demonstrate strong agreement with the main manuscript across the length spectrum. For the axonal volume, a significant negative gene-dosage effect was observed within shorter and medium-length tracts, mirroring the primary analysis. Consistent with the main findings, the myelin volume exhibited small effect sizes that did not reach statistical significance across any tract length. Finally, the aggregate g-ratio showed a robust, length-independent negative gene-dosage effect. This shift mirrors the primary manuscript, with deletion carriers demonstrating higher g-ratios and duplication carriers demonstrating lower g-ratios across the entire length spectrum.

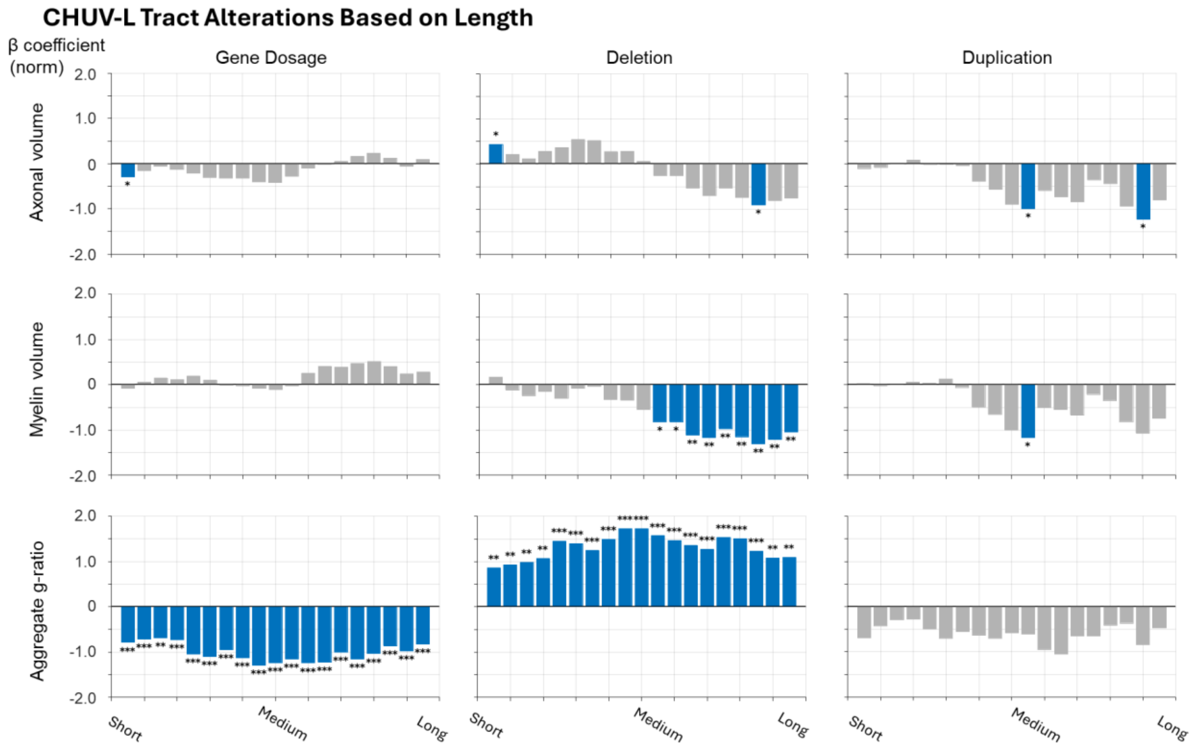


**Figure S6: Distribution of microstructural gene-dosage and categorical effect sizes across the tract-length spectrum in the CHUV-L dataset.** The x-axis represents white matter tracts ranked by physical length and grouped into 10% sliding-window bins. Bar heights represent standardized effect sizes (z-scores); gray bars indicate results that did not survive FDR correction. Findings for the axonal volume and g-ratio at this site closely align with the main manuscript. Specifically, the axonal volume exhibited a negative gene-dosage effect localized to shorter and medium-length tracts. For the g-ratio, a robust and length-independent negative gene-dosage effect was observed, consistent with the patterns reported in the primary analysis. However, results for the myelin volume at this site showed a deviation from the main manuscript; specifically, deletion carriers exhibited a significant negative effect within medium-to-longer tracts, whereas these effects were non-significant in the primary analysis. Despite this localized site-specific variation in myelin, the overall hierarchy of effects, defined by widespread g-ratio alterations and more localized axonal volume changes, remains highly consistent.

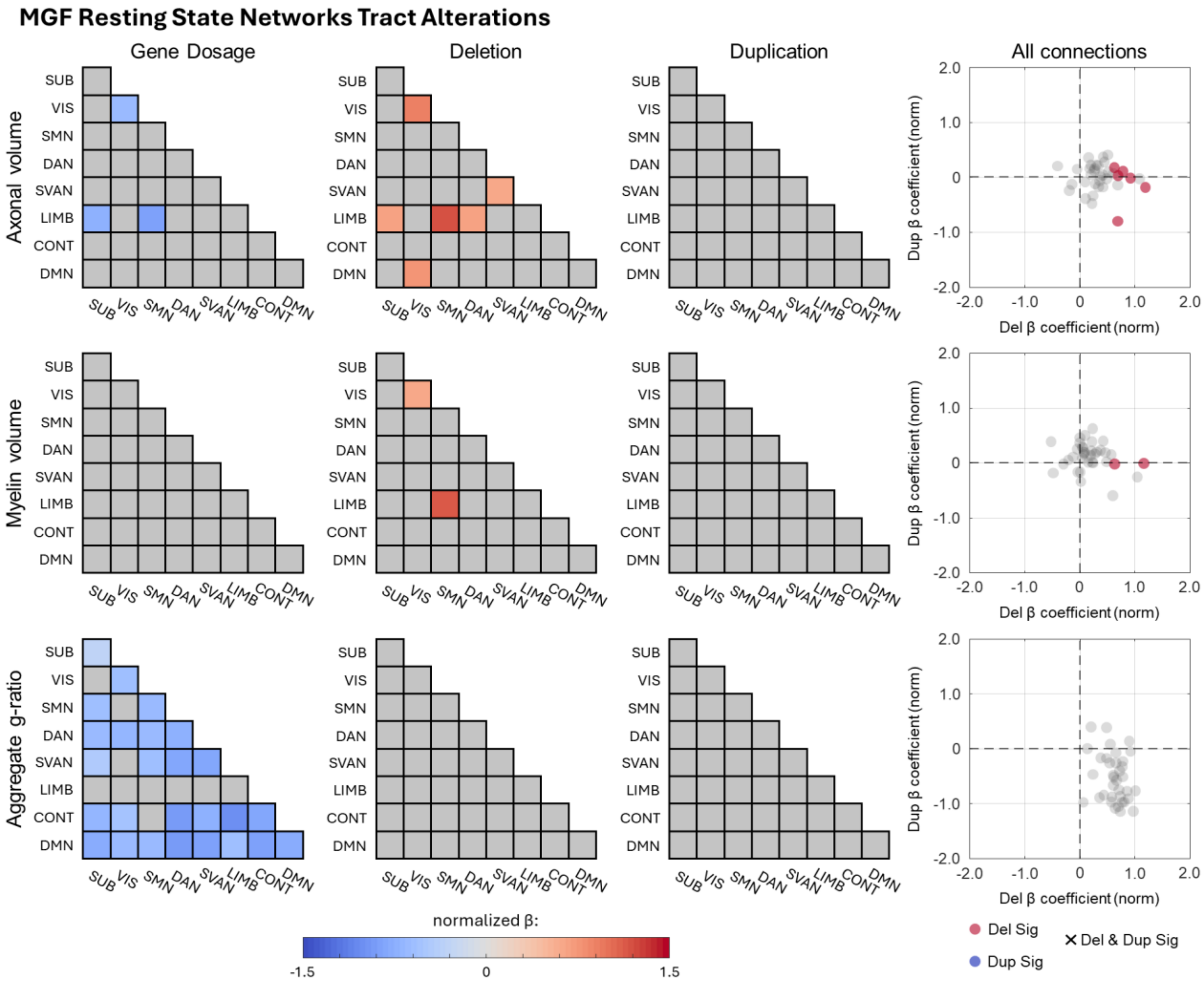


**Figure S7: Distribution of microstructural gene-dosage and categorical effects across functional networks in the MGF dataset.** The color scale represents standardized effect sizes (z-scores) relative to the site-specific normative baseline, with gray regions indicating functional connections that did not reach statistical significance following FDR correction. The rightmost column displays unthresholded effects to illustrate overall cohort trends regardless of significance. Consistent with the primary findings in the main manuscript, significant continuous gene-dosage effects were predominantly observed for both the axonal volume and the aggregate g-ratio, whereas the myelin volume exhibited no significant results. As illustrated by the unthresholded trends in the rightmost column, categorical analyses reveal that these overarching gene-dosage trajectories are driven primarily by alterations within the deletion cohort. Overall, the spatial distribution of these significant connections generally agrees with the network-level findings presented in the main manuscript. However, it is notable that while the continuous gene-dosage model for the g-ratio was significant, the corresponding isolated categorical models did not reach significance for either deletions or duplications at this specific site.

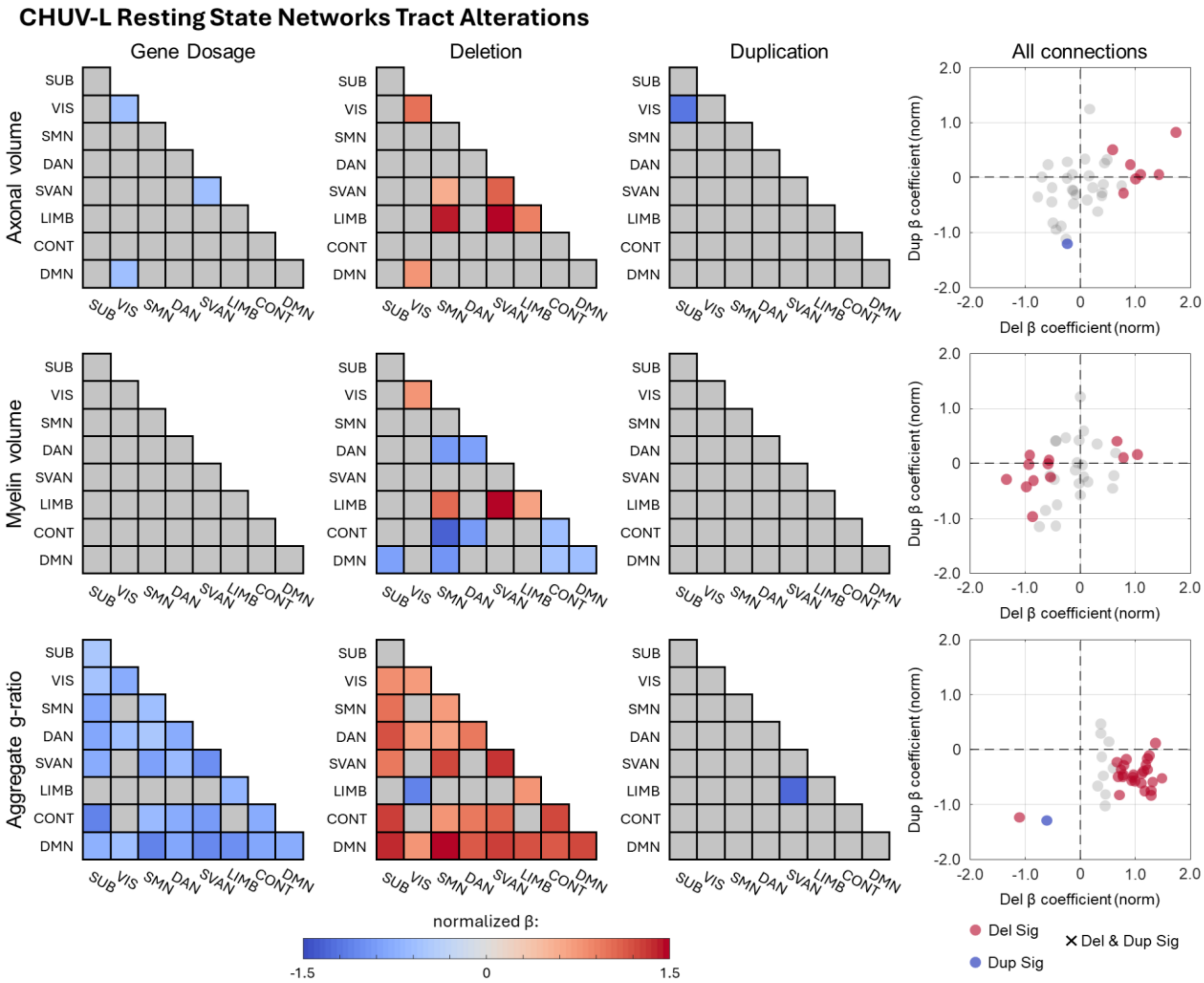


**Figure S8: Distribution of microstructural gene-dosage and categorical effects across functional networks in the CHUV-L dataset.** The color scale represents standardized effect sizes (-scores) relative to the site-specific normative baseline, with gray regions indicating nodes that did not reach statistical significance following false discovery rate (FDR) correction. The rightmost column summarizes categorical significance: red indicates nodes significantly altered in deletion carriers, and blue indicates significance in duplication carriers. Consistent with the primary findings in the main manuscript, significant continuous gene-dosage effects were observed for the axonal volume and the aggregate g-ratio. Categorical analyses revealed distinct spatial profiles at the nodal scale across these metrics. For the axonal volume, significant localized alterations were observed, a pattern driven primarily by the deletion cohort. In contrast, the myelin volume exhibited no significant continuous results and minimal nodal deviation, with only two isolated significant nodes identified in deletion carriers. This lack of significance aligns with the main manuscript's finding that 16p11.2 CNVs do not exert a strong or systematic unidirectional effect on myelin volume in either deletions or duplications. Finally, the aggregate g-ratio demonstrated the most widespread alterations across the network. As indicated by the categorical effects, this extensive nodal pattern is driven predominantly by deviations within the deletion cohort, mirroring the robust findings reported in the primary analysis.

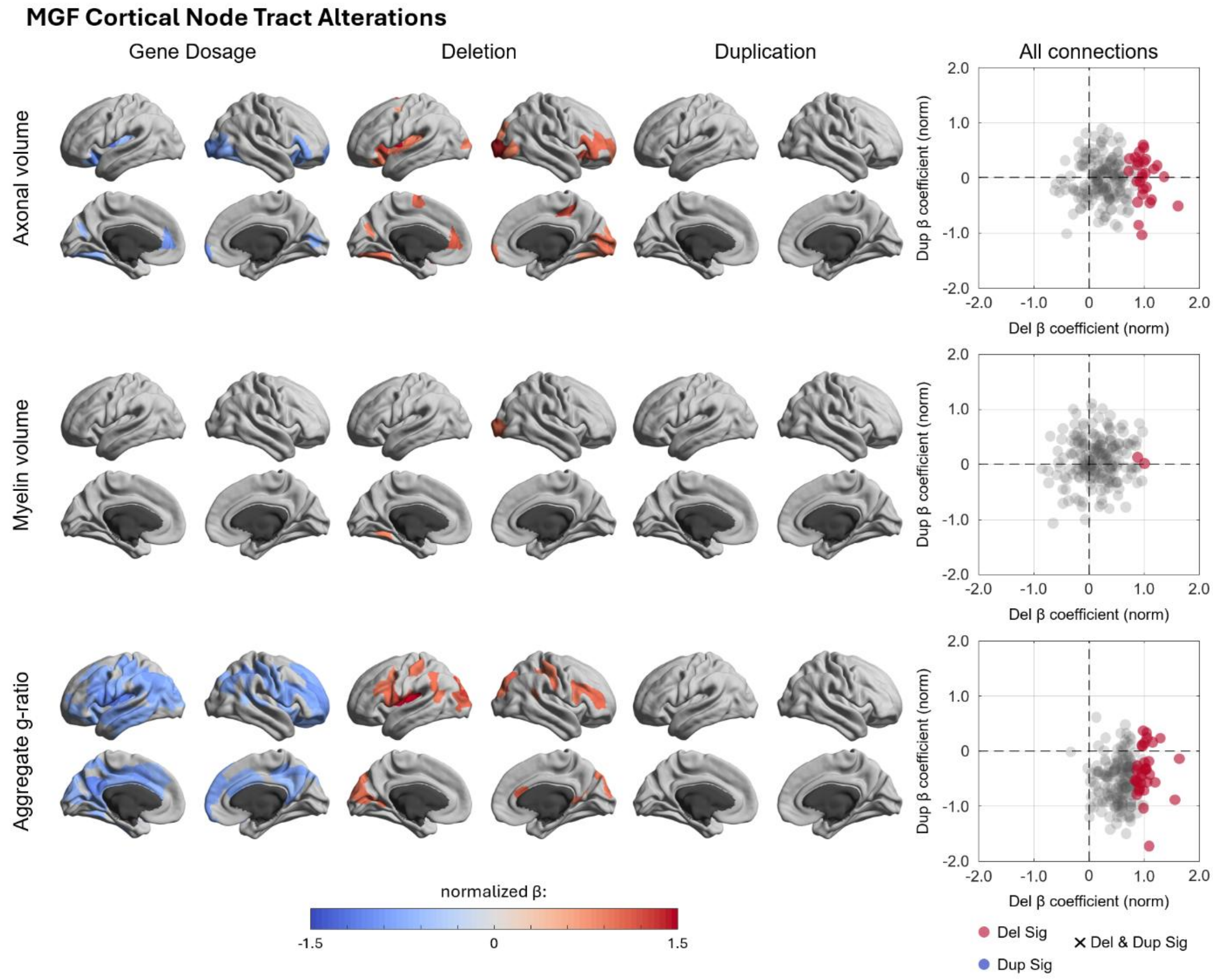


**Figure S9: Distribution of microstructural gene-dosage and categorical effects at the nodal scale in the MGF dataset.** The color scale represents standardized effect sizes (z-scores) relative to the site-specific normative baseline, with gray regions indicating nodes that did not reach statistical significance following false discovery rate (FDR) correction. The rightmost column summarizes categorical significance: red indicates nodes significantly altered in deletion carriers, blue indicates significance in duplication carriers. Consistent with the primary findings in the main manuscript, significant continuous gene-dosage effects were observed for the axonal volume and the aggregate g-ratio, whereas the myelin volume exhibited no significant continuous results. Categorical analyses revealed distinct spatial profiles at the nodal scale across these metrics. For the axonal volume, significant localized alterations were observed, a pattern driven primarily by the deletion cohort; a Wilcoxon test confirmed a significant difference in effect magnitudes between cohorts ($p < 0.001$). In contrast, the myelin volume exhibited no significant continuous results and minimal nodal deviation, with only two isolated significant nodes identified in deletion carriers. This lack of significance aligns with the main manuscript's finding that 16p11.2 CNVs do not exert a strong or systematic unidirectional effect on myelin volume in either deletions or duplications. Finally, the aggregate g-ratio exhibited the most widespread alterations across the network in the continuous gene-dosage model. While a Wilcoxon test comparing cohort effect magnitudes for the g-ratio did not reach statistical significance, the median absolute effect remained larger in the deletion cohort compared to duplications (0.62 vs. 0.52).

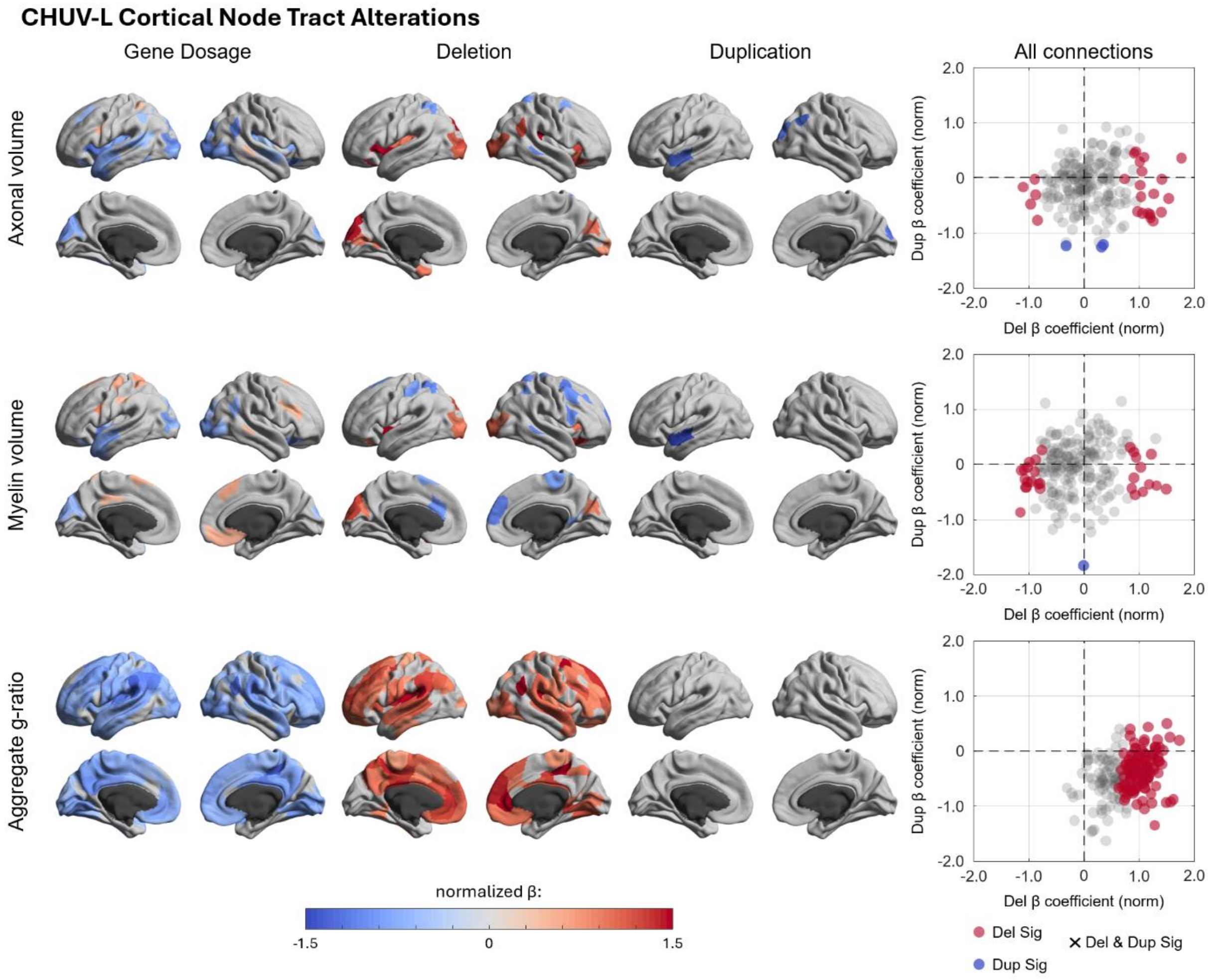


**Figure S10: Distribution of microstructural gene-dosage and categorical effects at the nodal scale in the MGF dataset.** The color scale represents standardized effect sizes (z-scores) relative to the site-specific normative baseline, with gray regions indicating nodes that did not reach statistical significance following false discovery rate (FDR) correction. The rightmost column summarizes categorical significance: red indicates nodes significantly altered in deletion carriers, and blue indicates significance in duplication carriers. Consistent with the primary findings in the main manuscript, significant continuous gene-dosage effects were observed for the axonal volume and the aggregate g-ratio. Categorical analyses revealed distinct spatial profiles at the nodal scale across these metrics. For the axonal volume, significant localized alterations were observed. To determine if this pattern was driven primarily by the deletion cohort, a Wilcoxon test on the absolute effect sizes was performed, confirming a significantly larger magnitude of alterations in deletion carriers compared to duplications ($p < 0.05$). In contrast, while the myelin volume exhibited a greater number of significant nodes in this cohort, these alterations were highly bidirectional. This spatial heterogeneity confirms the absence of a cohesive global trajectory, consistent with the main manuscript's observation that myelin volume alterations do not follow a systematic direction in either CNV group. Finally, the aggregate g-ratio exhibited the most widespread alterations across the network. To determine if this extensive nodal pattern was also driven predominantly by deviations within the deletion cohort, a Wilcoxon test on the absolute effect sizes was performed, confirming a highly significant difference in magnitude between the cohorts ($p < 0.001$). This finding mirrors the robust gene-dosage effects reported in the primary analysis.

### Methodological Validation and Comparison of Tract-Specific g-Ratio

Tract-specific g-ratio estimation represents a relatively novel framework for characterizing white matter microstructure. To validate the robustness of this approach, we benchmarked our results against state-of-the-art conventional tractometry. Theoretically, we anticipated a high degree of convergence in directional trends, where alterations in the g-ratio would be consistently reflected across both frameworks. The tractometry models utilized an identical two-step normative procedure to that of our tract-specific approach. However, total brain volume (TBV) was excluded as a covariate in the tractometry analysis. Because this methodology utilizes axonal volume fraction (AVF) and myelin volume fraction (MVF) rather than absolute volumes, the metrics are inherently scaled and spatially invariant; thus, they do not require adjustment for global head size.

Our results demonstrate broad agreement between the two methodologies across functional network and nodal scales. Both frameworks identified a significant negative gene-dosage effect across the spectrum, driven primarily by deviations in the deletion cohort. Notably, the tractometry approach yielded significant results across the entire deletion cohort. This strong effect likely reflects the influence of partial volume effects, a phenomenon in whole-brain voxel-based metrics that may increase sensitivity to global alterations at the cost of specificity.

In the analysis of physical tract lengths, we observed a localized divergence in the MVF results. Specifically, the tractometry approach revealed a significant positive effect in duplication carriers for medium-to-long tracts, which subsequently manifested as a significant continuous gene-dosage effect. It is important to note, however, that longer connections traverse a greater number of voxels, inherently increasing their susceptibility to partial volume averaging relative to shorter tracts. Furthermore, the p-values for these MVF findings were substantially higher than the effects observed for AVF and the g-ratio. Ultimately, when evaluating the tractometry results, the microstructural phenotype is characterized by a negative gene-dosage effect in both AVF and the g-ratio, driven predominantly by deletion carriers with minimal impact on myelin, which demonstrates strong agreements with the findings obtained from our primary tract-specific framework.

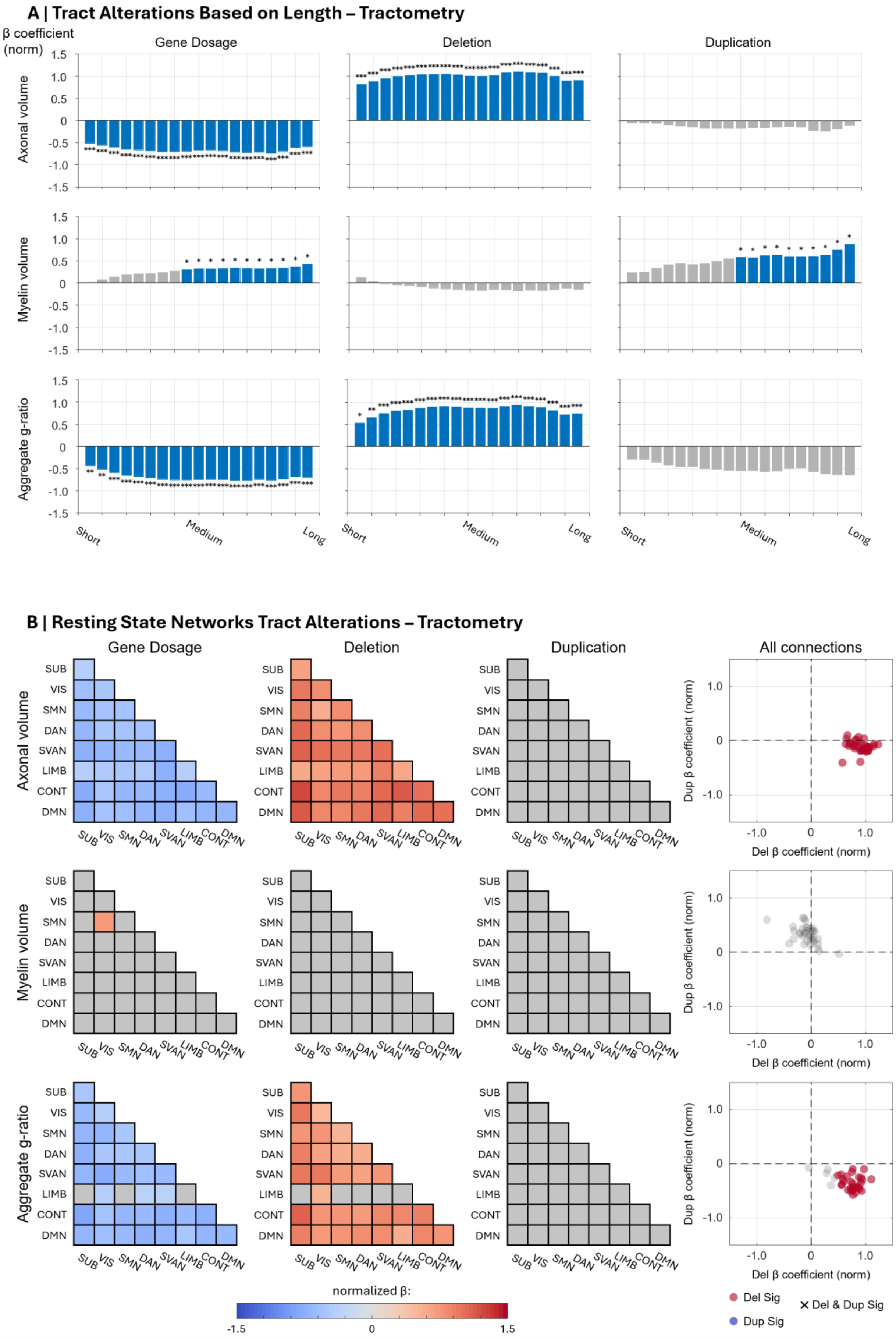
A | Tract Alterations Based on Length – Tractometry
β coefficient (norm)
Gene Dosage
Deletion
Duplication
Axonal volume
Myelin volume
Aggregate g-ratio
Short
Medium
Long
B | Resting State Networks Tract Alterations – Tractometry
Gene Dosage
Deletion
Duplication
All connections
SUB
VIS
SMN
DAN
SVAN
LIMB
CONT
DMN
Dup β coefficient (norm)
Del β coefficient (norm)
normalized β:
-1.5
0
1.5
Del Sig
Dup Sig
Del & Dup Sig

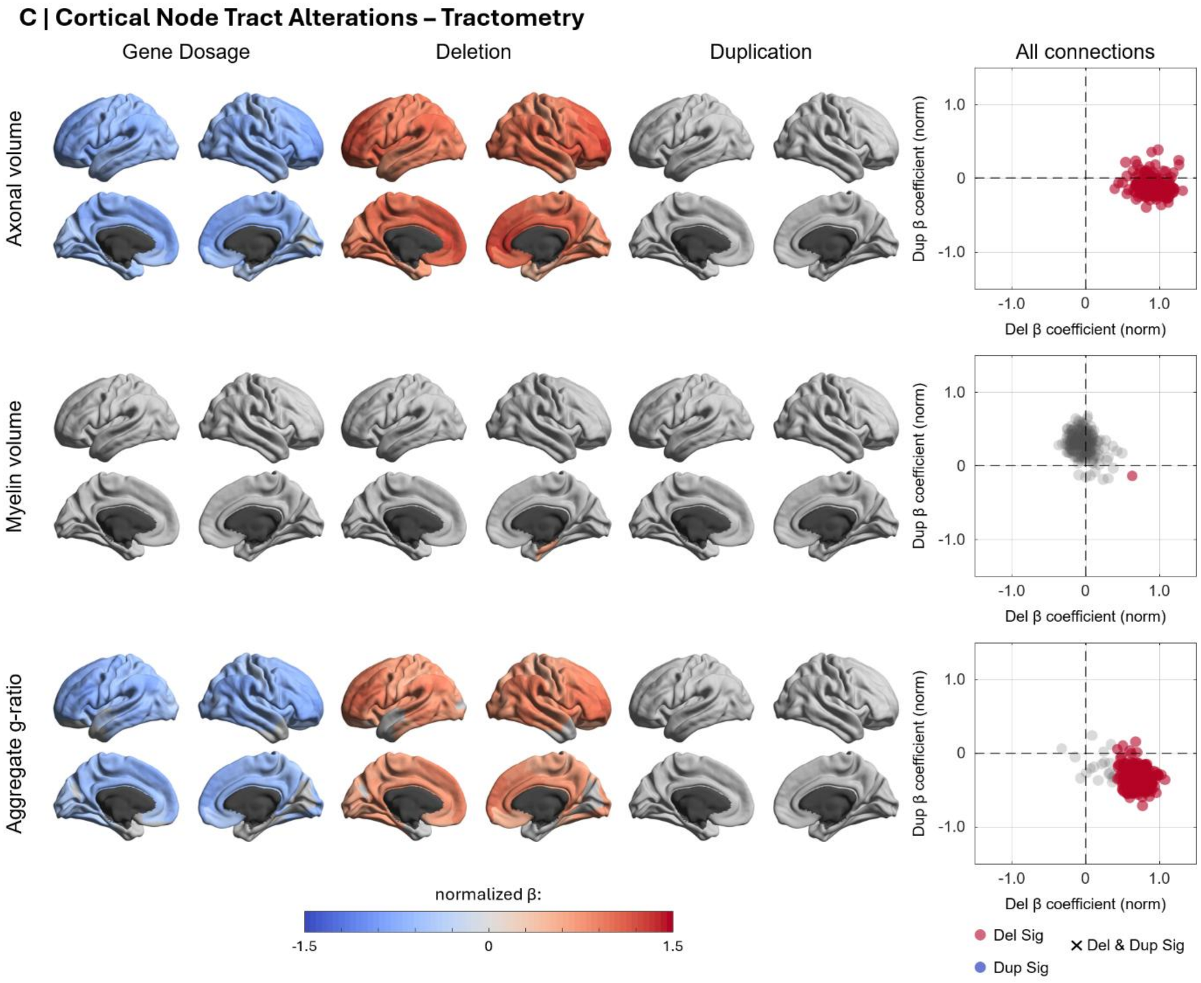


**Figure S11: Cross-scale validation of microstructural gene-dosage and categorical effects using tractometry.** This figure benchmarks the primary results against state-of-the-art conventional tractometry across the tract-length spectrum (A), functional networks (B), and cortical nodes (C). The color scale represents standardized effect sizes (z-scores) relative to the normative baseline, with gray regions indicating results that did not reach statistical significance following FDR correction. Consistent with the primary findings, significant continuous gene-dosage effects were observed for the AVF and the aggregate g-ratio across all scales, whereas the MVF generally exhibited no significant continuous results. These overarching trajectories were driven primarily by alterations within the deletion cohort. For the tract-length analysis, findings for the AVF and the aggregate g-ratio demonstrate strong agreements with the primary tract-specific framework. However, tractometry approach identified significant positive MVF deviations in duplication carriers for medium-to-long tracts, these results warrant caution; longer connections exhibit a greater susceptibility to partial volume averaging, and the associated p-values were higher than those observed for the AVF and g-ratio results. At the nodal scale, the tractometry approach yielded significant results across the entire deletion cohort, likely reflecting the increased sensitivity of whole-brain voxel-based metrics to global partial volume effects
